# A rigorous framework for detecting SARS-CoV-2 spike protein mutational ensemble from genomic and structural features

**DOI:** 10.1101/2021.02.17.431625

**Authors:** Saman Fatihi, Surabhi Rathore, Ankit K. Pathak, Deepanshi Gahlot, Mitali Mukerji, Nidhi Jatana, Lipi Thukral

**Affiliations:** CSIR-Institute of Genomics and Integrative Biology, Mathura Road, New Delhi, 110 025, India; Academy of Scientific and Innovative Research (AcSIR), CSIR-Human Resource Development Centre, (CSIR-HRDC), Kamla Nehru Nagar, Ghaziabad, 201002, Uttar Pradesh, India

**Keywords:** SARS-CoV-2, spike protein, mutations, lineages, structural features

## Abstract

The recent release of SARS-CoV-2 genomic data from several countries has provided clues into the potential antigenic drift of the coronavirus population. In particular, the genomic instability observed in the spike protein necessitates immediate action and further exploration in the context of viral-host interactions. By temporally tracking 527,988 SARS-CoV-2 genomes, we identified invariant and hypervariable regions within the spike protein. We evaluated combination of mutations from SARS-CoV-2 lineages and found that maximum number of lineage-defining mutations were present in the N-terminal domain (NTD). Based on mutant 3D-structural models of known Variants of Concern (VOCs), we found that structural properties such as accessibility, secondary structural type, and intra-protein interactions at local mutation sites are greatly altered. Further, we observed significant differences between intra-protein networks of wild-type and Delta mutant, with the latter showing dense intra-protein contacts. Extensive molecular dynamics simulations of D614G mutant spike structure with hACE2 further revealed dynamic features with 47.7% of mutations mapping on flexible regions of spike protein. Thus, we propose that significant changes within spike protein structure have occurred that may impact SARS-CoV-2 pathogenesis, and repositioning of vaccine candidates is required to contain the spread of COVID-19 pathogen.

## 1. Introduction

The current novel coronavirus disease 2019 (COVID-19) outbreak is caused by severe acute respiratory syndrome coronavirus 2 (SARS-CoV-2) (Andersen et al., 2020). By August 2021, the World Health Organisation (WHO) reported 207,784,507 confirmed cases with 4,370,424 deaths all over the world. Global sequencing initiatives have been indispensable in understanding SARS-CoV-2 genomic diversity. There is a great deal of interest in identifying how spike protein of SARS-CoV-2 is evolving as it directly interacts with the human ACE2 (hACE2) receptor (Walls et al., 2020). Efforts to understand mutations and predict associated pathogenesis have been hampered by our inability to evaluate combinations of mutations.

The spike protein has multidomain structural architecture and comprises S1 and S2 subunits (Walls et al., 2020). The viral infection commences upon binding of the S1 subunit to the hACE2, followed by the proteolytic cleavage in the spike protein resulting in the dissociation of the S1 and S2 subunits, further leading to membrane fusion and virus entry inside the cell (Hoffmann et al., 2020b). S1 subunit is the key domain responsible for mediating the receptor binding and hence is considered to be the potential target for vaccines and drugs against COVID-19 (Lan et al., 2020; Dai and Gao, 2021; Wang et al., 2021). S1 subunit (1-685 a.a.) is comprised of a signal peptide followed by the N-terminal domain (NTD), and the receptor-binding domain (RBD) (Walls et al., 2020). Although the function of the NTD is not clearly known, it has been suggested to harbor the neutralizing epitopes (Chi et al., 2020; Liu et al., 2020; Graham et al., 2021). The RBD is a well-established domain that is characterized into a receptor-binding motif (RBM) and the core region (Mittal et al., 2020; Shang et al., 2020). Residues of the RBM play an important role in mediating key interactions with the hACE2 receptor (Lan et al., 2020). Many RBD targeting antibodies interfere either with RBD-hACE2 binding or inhibits the RBD to acquire the up conformation (Chen et al., 2020; Barnes et al., 2020; Ju et al., 2020). Recent studies on monoclonal antibody response against mRNA vaccines have shown a codominant antibody binding to the NTD and RBD, thereby making NTD an equally important therapeutic target (Amanat et al., 2021). Another interesting functional motif is the connecting linker region between the RBD and HR1 that contains two critical furin cleavage sites, namely- S1/S2 site and S2′ followed by a fusion peptide (Xia et al., 2020; Hoffmann et al., 2020a). The understanding of these regions and how they are evolving during the current pandemic is critical for enhancing present vaccine efficacy.

Early analysis of viral variants revealed a single amino acid substitution D614G, which has now become a widespread dominant variant (Korber et al., 2020). Although it did not reveal greater clinical severity, structural features of mutant D614G spike showed interesting characteristics (Volz et al., 2021; Groves et al., 2021). The conformational state of spike protein was found more stable in an “up” conformation (Yurkovetskiy et al., 2020). In Oct 2020, multiple co-occurring mutations in the SARS-CoV-2 within spike protein were reported (Rees-Spear et al., 2021). Three major lineages, including B.1.1.7 (Alpha), B.1.351 (Beta), and P.1 (Gamma) were seen emerging in patients from the UK, South Africa, and Brazil, respectively (Rambaut et al., 2020; Tegally et al., 2020; Resende et al., 2021), which are now classified as the VOCs. Compared to previous variants, the Alpha variant raised concern due to the presence of N501Y mutation in the RBD of the spike protein. Particularly in spike, this variant emerged with several other mutations like 69/70 deletion and P681H mutation present in the N-terminal domain (NTD) and near S1/S2 site, respectively (Rambaut et al., 2020). Another variant from South Africa (B.1.351) emerged independently with multiple mutations in the RBD (K417N and E484K) along with N501Y, however, lacked the 69/70 deletion mutations in the NTD segment (Tegally et al., 2021). The lineage B.1.526 is classified as a Variant of Interest (VOI) which have multiple spike mutations with unique substitutions L5F, T95I, and, D253G, which were not present earlier in any other lineages (Annavajhala et al., 2021; West Jr et al., 2021). Recently, a rapidly transmitting variant emerged in India, namely-B.1.617.2 (Delta) with unique RBD mutations E484Q and L452R, two deletions (E156del and F157del) and a substitution (R158G) in the NTD and 4 other mutations including D614G (Kannan et al., 2021). A sub-lineage of Delta with an additional mutation, K417N is termed as AY.1 ( Delta-plus). Interestingly, each lineage contains more than a handful of spike mutations also present on non-RBD regions. Although they are not the focus of extensive characterisation, they are likely to contribute in a specific manner towards spike evolution. A better structural understanding of pattern underlying these lineages, as a whole will facilitate in predicting the immune response with new SARS-CoV-2 variants.

In this work, we integrated large-scale genomic variations of SARS-CoV-2 spike protein with extensive structural characterisation. We identified mutational hotspots based on a temporal acquisition of mutations and found distinct variant and invariant patches distributed throughout the structure. The contribution of lineage-defining mutations was evaluated domain-wise to understand structural motifs that are accumulating more mutations than others. Some of these mutations have also evolved into VOCs. Based on mutated structural models of known VOCs, we found that structural properties such as accessibility, secondary structural type, and intra-protein interactions at local mutation sites are greatly altered. Extensive molecular dynamics simulations of D614G mutant spike structure with hACE2 further revealed dynamic features that may play a key role in the conformational ensemble of the spike protein. Our results support the view that structural changes of prominent SARS-CoV-2 lineages significantly alters structural properties of spike protein and hence may require updating spike-based vaccine candidates.

## Results

### Genomic diversity of spike protein

We present meta-analysis of SARS-CoV-2 genomic data from GISAID sampled until May 2021. Amino acid substitutions, including missense and deletions/insertions were extracted for the spike protein. In total, 6,517 mutations were observed in 1,624,465 spike sequences. The frequency of each substitution was mapped onto the full-length spike structure model as shown in (Figure 1). Both S1 and S2 subunits are marked, with the S1 domain of the SARS-CoV-2 consisting of the N-terminal domain (NTD: 14-305 a.a.) and the receptor binding domain (RBD: 319-541 a.a.) clearly accumulating more mutations. The S1 subunit harbours the prominent D614G mutation present in 99.84% of genomes (Isabel et al., 2020). The sequence alignment conservation with spike protein homologs also revealed that the S1 subunit is highly variable compared to the S2 stalk, with the HR1 region as the most conserved segment (Figure S1). While majority of the residues show low-frequency mutation rate, two major frequency groups were observed. Eight prominent mutations were present in the first group with more than 50% frequency; N501Y, P681H, T716I, D1118H, A570D, S982A, HV69/70del and Y144del. The second group with frequency between 1-20% displayed 32 unique residues, with 53% and 18% amino acids present in the NTD and RBD domains, respectively. We thus, were able to associate mutations with structural domains and next utilized large-scale genomic data to understand domain-wise temporal acquisition of mutations.

**Figure 1:**
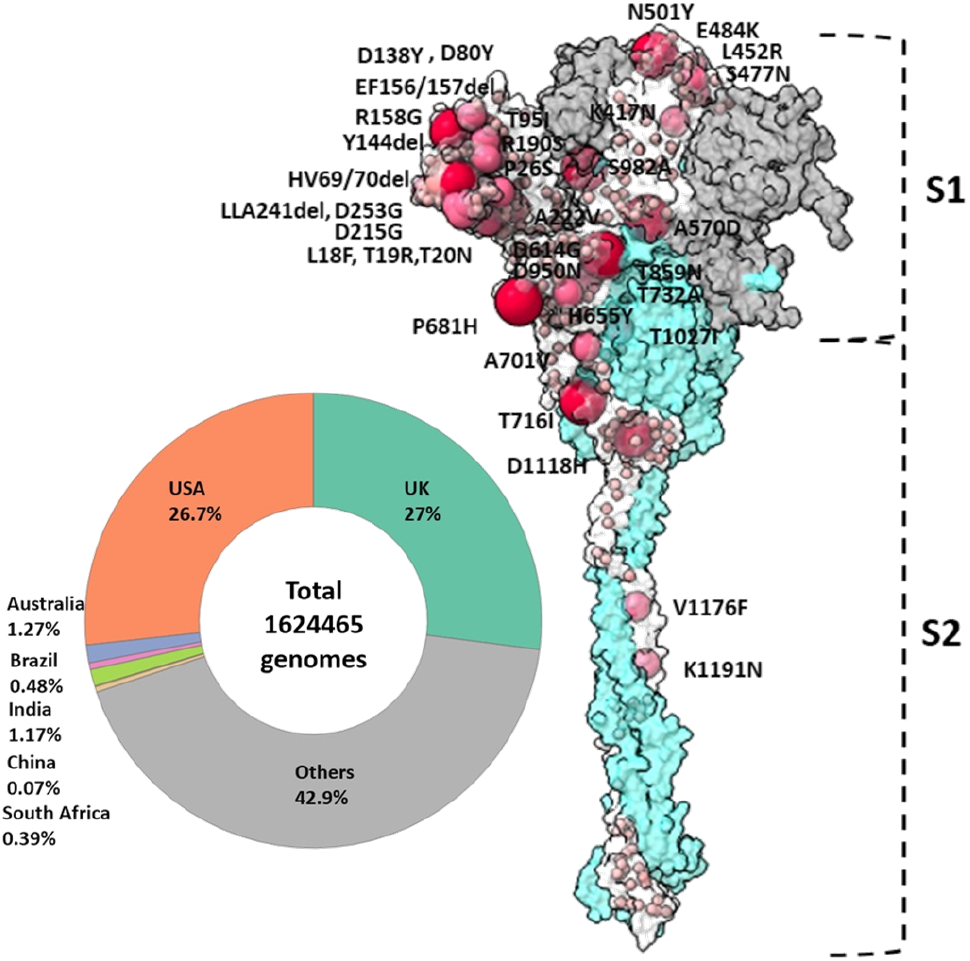
Genomic Diversity of SARS-CoV-2 spike protein. In total, 1,624,465 genomes were retrieved from GISAID, as plotted in the pie chart, with different colors showing country-wise submitted genomes till May 2021. The modelled full-length spike protein is shown in surface representation, with the S1 and S2 subunit colored in grey and cyan, respectively. In total, 6,517 mutations were mapped on one protomer, out of which 1,312 were found to be on unique sites. The size of the sphere indicates frequency of mutations, with dark to low gradient showing a higher to lower frequency range. The amino acids mutating at higher frequency are marked in text.

### Temporal analysis of mutational hotspots reveals hypervariable regions on spike protein

How did distinct protein regions evolve since the start of the covid-19 pandemic? To answer this, we located high-frequency mutations (>1% frequency) on the spike structure to understand the invariant and variant patches on different spike domains. Not all residue positions are susceptible to mutations. A clear trend emerging from dynamic tracking of genomic data is the constant evolution of specific S1 domains in the NTD, RBD, and linker regions (Figure 2A; Figure S2). In early pandemic, L5F substitution in the signal peptide was present, alongwith D614G mutation in the linker. From July 2020, noticeable accumulation of mutations were observed in the NTD and RBD regions. The S2 stalk showed least number of mutations, with three variable segments, specifically the L1 region (701-716 a.a.), S982 residue in the HR1, and D1118 residue in the loop region connecting HR1 and HR2 region.

**Figure 2:**
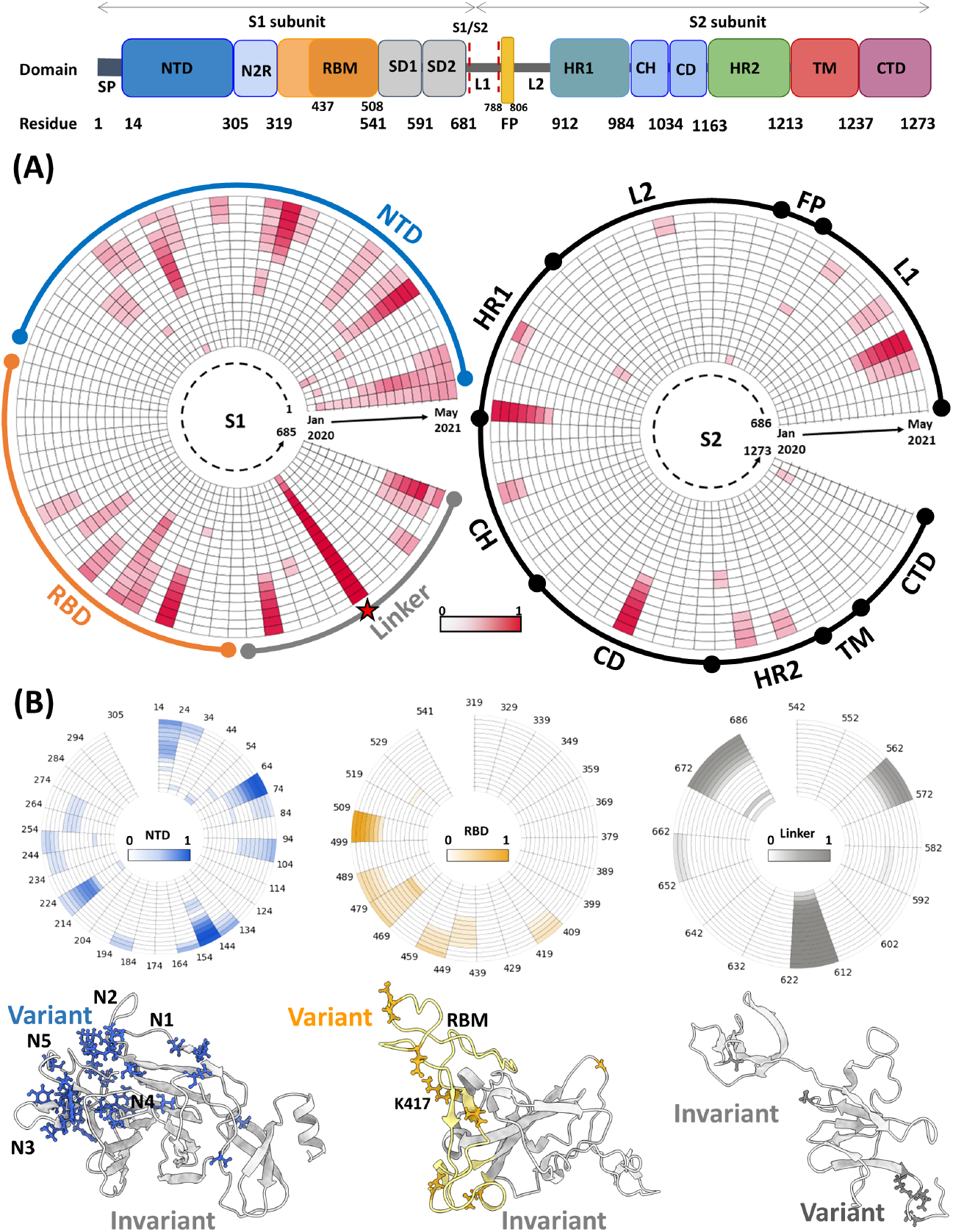
Mutation clusters of SARS-CoV-2 spike protein. (A) The schematic view of the domain architecture of SARS-CoV-2 spike protein is shown in the top row. The circular plot shows the ‘normalised mutation frequency’ in a time-dependent manner. Each row represents one month and, in total 17-month period from January 2020 to May 2021 is shown. The spike protein of 1273 residues is divided into 129 ten-residue bins and is shown for S1 and S2 subunits separately. The normalised mutation frequency is calculated as the ratio of *N*_*i*_ and N_ref_. The *N*_*i*_ refers to cumulative number of mutations from global GISAID data for each bin and N_ref_ is the bin that corresponds to maximum frequency bin. The color gradient of red denotes the intensity of mutation frequency, with 1 being the highest. The D614G present in the linker region is marked as a star for reference as it is now present in ≈98% genomes.(B) Zoomed in circular plots are shown for three major parts of the S1 subunit; NTD, RBD and Linker regions. The corresponding structural mapping for each domain is shown to depict mutational spots. The mutation site is depicted in stick representation highlighting the variant patches.

To look at deeper level, a quantitative description of the time progression of highly variable S1 domain regions is shown in Figure 2B. Within the NTD, five prominent variant stretches were observed, with Y144 deletion, G142D, E156-F157 deletion, and R158G segment as the largest contiguous patch. The N-terminal region of the NTD, namely L18F, T19R, T20N, evolved early during the pandemic. Later during the pandemic, several segments in N1, N3, and N5 loops of NTD mutated significantly faster, indicating that these regions are hypervariable segments, also shown as protein snapshots. Interestingly, the connecting region between the NTD and RBD (N2R: 310-319) is not mutating. While the core of the RBD consisting of five stranded antiparallel *β*-sheet is an invariant region, the RBM motif that binds to the hACE2 receptor harbored 99% of the RBD mutations. Only a single K417 residue is present outside the RBM, however, it is still at the interface and is known to form a salt bridge with D30 of the hACE2 receptor. The linker region consists of variant stretches contributed from three prominent mutations A570D, H655Y and P681H/R along with D614G. Overall, spike protein harbors multiple prevalent mutations which form mutational clusters that have relatively evolved more than others. Apart from the well-characterized functional significance of RBM, other hypervariable mutational clusters have relatively less functional data. We speculate that these protein regions are important for the function or stability of protein and hence are under greater selective pressure.

### Integrating defining spike mutations within lineages reveal coaxiality at NTD

SARS-CoV-2 is continually evolving and population genomics studies reveal constellation of mutations that define a particular lineage. In principle, lineages reveal pool of mutations that is characteristic of subset of viral population. To gather insights on co-occurring mutations, we obtained lineage information of SARS-CoV-2 from PANGOLIN database. Figure 3A shows the time evolution of 43 SARS-CoV-2 lineages. Strikingly, number of spike mutations have significantly increased since evolution of B.1.1.7 lineage in October 2020 (Figure S3). For instance, the recent Delta lineage accumulated a total of 9 defining spike mutations throughout the protein.

**Figure 3:**
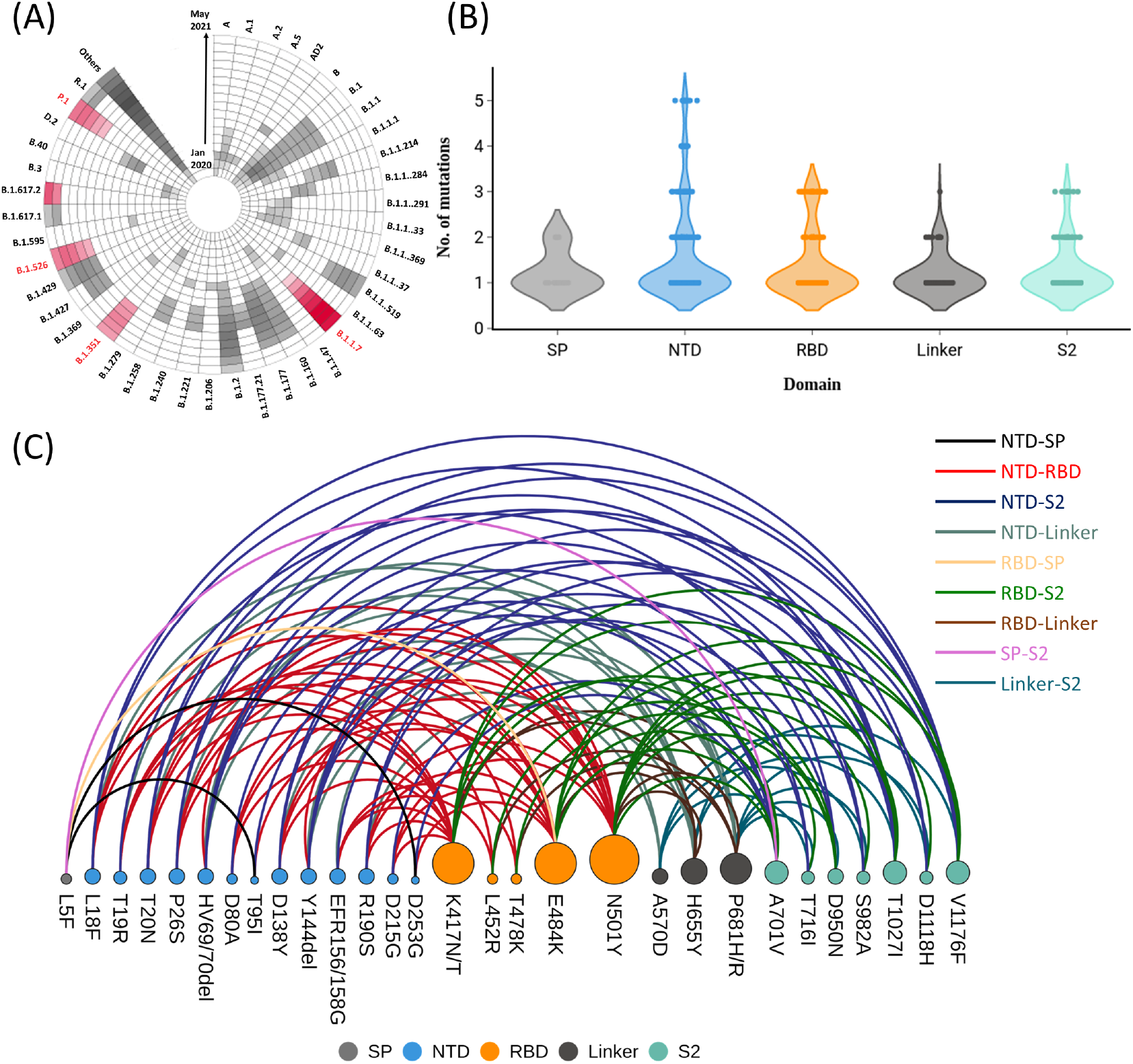
Dynamical tracking of genomes reveal spike protein mutational ensemble. (A) The circular plot shows temporal progression of occurrence of top lineages over a span of 17 months (January 2020-May 2021). The color gradient highlights the prevalence of particular lineage in each month, where the key lineages are highlighted in red color. (B) The violin plot depicts domain wise distribution of defining mutation present in each lineage where each dot represents a lineage. (C) The arc diagram plots the network of co-occurring mutations between different spike domains, where the size of the node denotes the degree and edges are colored according to the inter-domain connections.

The lineage data also allowed us to examine the underlying link with combination of mutations and structural domains. For example, are these co-occurring mutations emerging in a specific pattern? We identified that lineage-defining mutations were maximally located on the NTD region followed by RBD, Linker, S2 subunit, and signal peptide (Figure 3B). In particular, 5 NTD residues appeared in combination that persisted in multiple lineages. On the other hand, fewer RBD residues were involved. A deeper understanding of residue-wise distribution showed that specific domain combinations were preserved in six major lineages, including five VOCs. We observed that majority of the combinations occur within four domains, with NTD as a key co-occurring domain (Figure 3C). For instance, L18F, T20N, P26S, D138Y and R190S are present in P.1 lineage. Similarly, the recent B.1.617.2 lineage possess 4 NTD mutation combination (T19R, E156del, F157del, and R158G). The RBD residues, on the other hand, utilise less number of mutations that have overlapped with number of lineages. A single amino acid substitution L452R is present in multiple lineages, including B.1.526, B.1.427, B.1.429, B.1.617.1, B.1.617.3, and lineages designated as Delta (B.1.617.2, AY.1, AY.2, and AY.3). Similarly, E484K is present in B.1.525 (Eta), P.1 (Gamma), and B.1.351 (Beta). The combination of K417N, E484K, and N501Y substitutions were also present in B.1.351 (Beta) and P.1 (Gamma). The linker region with P681R and H655Y along with D614G were the main amino acid co-occurring substitutions. In addition, NTD also has more intra-domain mutation combinations across VOC linegaes.

Overall, there exists a pattern of mutational clusters with highest fraction of inter-domain combinations mapping to the NTD-RBD, NTD-linker and NTD-S2 regions. These results indicate that NTD forms a coaxial cluster that is under stronger selective pressure than other domains.

### Structural determinants of mutations in VOCs reveal higher accessibility and disordered loops

Are these co-occurring mutations altering the spike topology or its interface that may impact versatility of protein interactions? In order to understand molecular level details, we generated six glycosylated spike protein models for key lineages (Figure 4A). These structures now provide us with spatial maps that include amino acid substitutions and in some cases deleted segments. Most of these changes render the local region with an increased surface accessibility (Figure 4B). We analysed the change in the relative surface area (RSA) for each characteristic mutations in these lineages. In total, ≈70% of the characteristic mutations in B.1.351 and Delta lineages showed a significant increase in the surface area, and the rest remained unchanged. The P.1 and B.1.526 lineages showed limited increased RSA for 40% and 16.7% of its defining mutations, respectively. Interestingly, the average RSA for the entire RBM across all the key lineages (39.05 Å^2^) was comparable to WT RBM (39.75 Å^2^), except for the B.1.526 lineage which displayed a relatively lower average RSA (37.44 Å^2^) for its RBM region.

**Figure 4:**
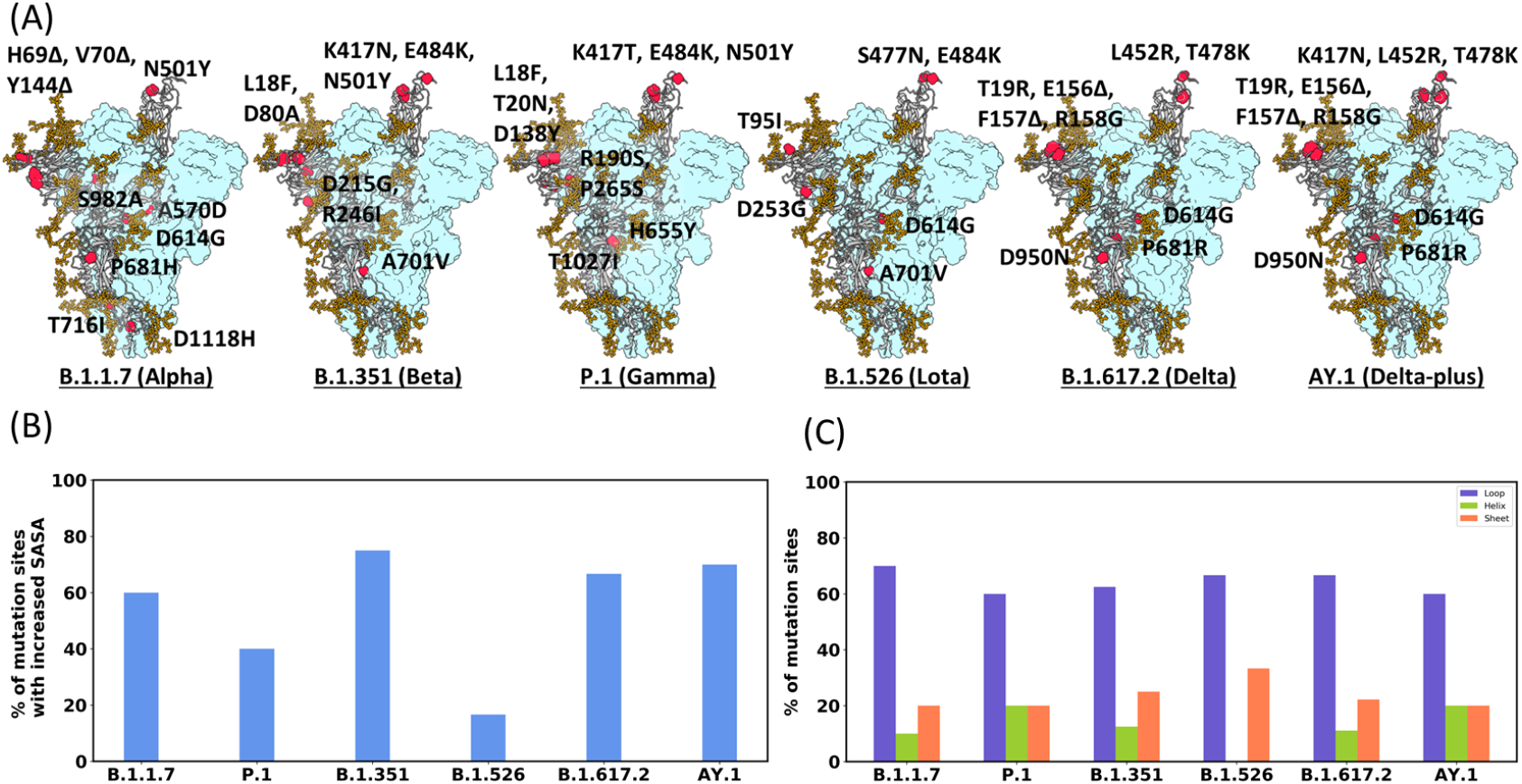
Altered relative surface area and presence of the mutations on variable loops. (A) Structural snapshots of the characteristic mutations in each key lineage are highlighted in red spheres on one of the chains of the spike trimer, with the glycans highlighted in brown. (B) The relative surface area for each mutation site in all variant models were analysed. The bar plot depicts the percentage of the characteristic mutations resulting in an increased surface accessibility as compared to the WT. (C) The grouped bar plot shows the quantitative measure for the secondary structure assessment of each variant and the presence of their respective characteristic mutations on the variable loop region, sheet, and helix of the spike protein.

Next, we investigated the secondary structural features of the regions where these substitutions and deletions occur in these key lineages (Figure S4). Interestingly, majority of the characteristic mutations (21 out of 35) in these selected key lineages reside within variable loop regions, as can be seen from Figure 4C. 70% of the characteristic mutations in the B.1.1.7 lineage occur in the loop regions. Similarly, in the B.1.526 and B.1.617.2 lineages, 66.7% of the characteristic mutations are harbored in the loops, whereas in P.1, B.1.351, and AY.1 lineages, the location of these mutation sites quantifies to be 60-62.5%. Further, 12 out of 15 mutations in the NTD and 4 out of 7 mutations in the RBD across all five VOCs and B.1.526 fall in the variable loop region. These findings reveal that mutations are relatively increasing the surface accessibility of mutated loci and loop regions are accruing more mutations.

### Probing local interaction network of spike protein revealed dense hotspots in Delta variant

We were further interested in understanding how combination of amino-acid substitutions may alter overall protein interaction network. By comprehensively analysing the intra-molecular hydrogen bonds, salt-bridges and hydrophobic contacts in these key lineage structural models, we observed significant variations in non-covalent bond pattern compared to wild-type structure (Figure 5; Figure S5). Number of hydrogen bonds increased by 34.8% in these variants as compared to the WT with 330 unique hydrogen bonds formed. For instance, in B.1.617.2 (Delta variant) four hydrogen bonds upon T19R mutation were found, whereas only one hydrogen bond was observed at this site in the WT protein structure. Whereas, one of the two major deletions in the NTD of B.1.617.2 variant resulted in the loss of 4 hydrogen bonds mediated by E156 in the WT (Figure S6). Another consecutive substitution, R158G also perturbed the formation of 5 hydrogen bonds in the variant. The RBD of B.1.617.2 harbors two defining mutations (L452R and T478K), out of which L452R substitution enhanced the local hydrogen bond networking by forming 3 bonds as compared to the one in WT (Figure 5B; Figure S7). There were 18 salt-bridges in the WT spike which were found to be retained in the VOCs. However, 8 unique salt-bridges were observed in the variants, out of which 2 lie in the NTD, 2 in the RBD and 4 in the S2 subunit. The number of interactions driven by hydrophobic residues also significantly increased, with 32 novel contacts observed in the VOCs.

**Figure 5:**
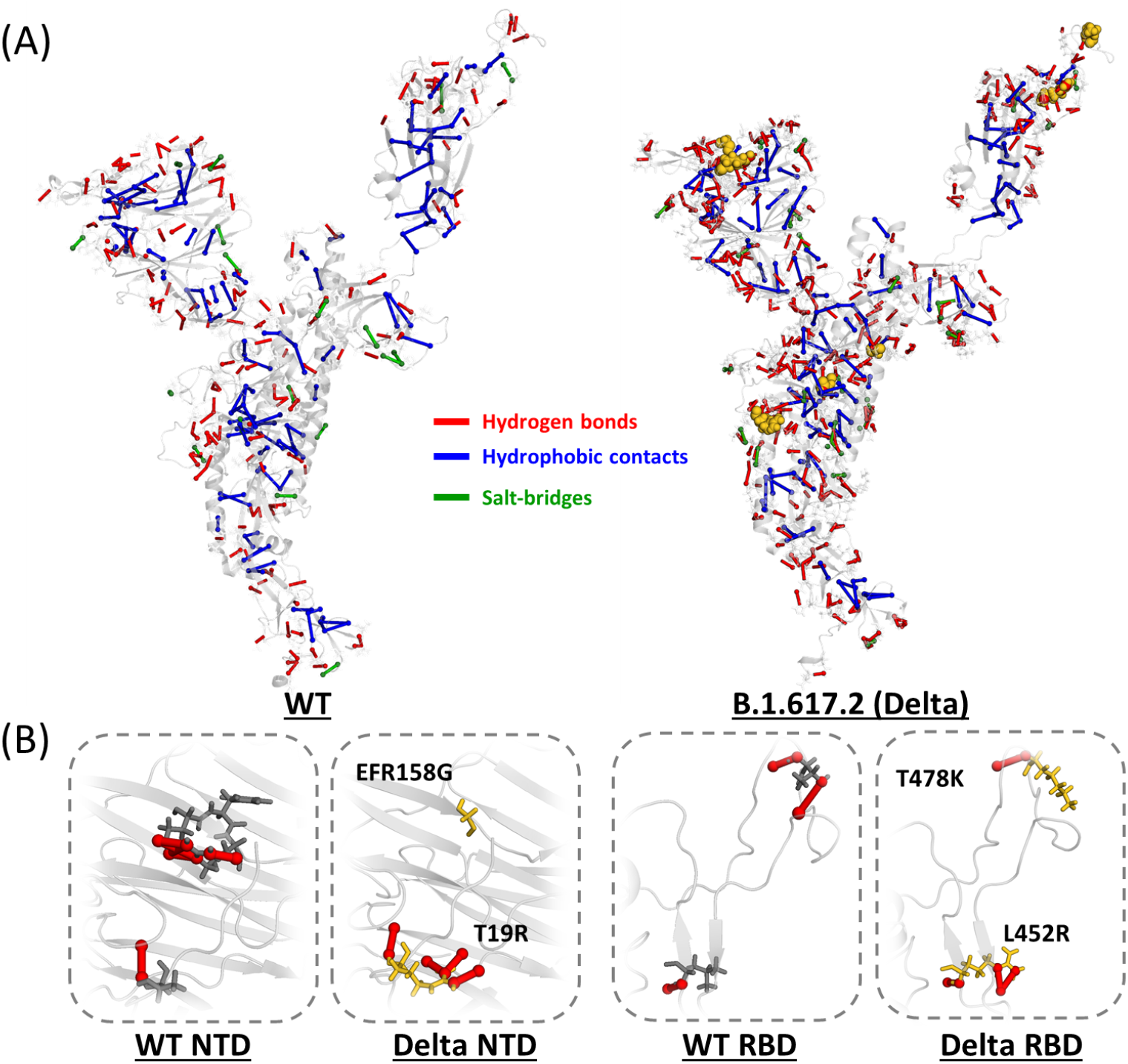
Intra-protein interaction network of WT vs B.1.617.2 spike. (A) Comparative global intra-molecular network for WT and Delta spike protein, where unique hydrogen bonds are marked in red on one of the chains of spike trimer. The total number of salt bridges and hydrophobic contacts are highlighted in green and blue, respectively and the mutations of B.1.617.2 are marked in yellow. (B) The zoomed-in panels display the comparison between local hydrogen bond network around the characteristic mutations for which the side chains are shown in stick representation in the NTD and RBD of the WT and Delta variant.

Next, to examine the impact of these mutations on the protein structure, we calculated domain-wise residue interaction network (RIN) in wild-type and six key spike variants, namely- B.1.1.7, B.1.351, P.1, B.1.526, B.1.617.2, and AY.1. In these networks, residue represents node and contacts between them are edges. Comparative analysis of networks of VOCs with wild-type revealed perturbations in local contacts and there is an overall increase in the total number of intra-protein contacts in these key lineages. Remarkably, the intra-molecular RBD contacts increased significantly with 660 additional contacts or edges and 70% of the residues displayed a degree >10. In B.1.1.7 lineage, there is one critical mutation in the RBM namely- N501Y, which has been reported previously to enhance the interaction with the hACE2 (Zahradnik et al., 2021; Zhu et al., 2021). Hence, we analyzed the inter-protein interaction network of the binding interface between the RBD and hACE2 (Figure S8). Structural analysis of the WT and B.1.1.7 variant bound to hACE2 revealed the formation of a high number of contacts after N501Y substitution. Similarly, P.1 and B.1.351 lineages have three key mutations in the RBM, namely-K417T/N, E484K, and N501Y. For K417T in B.1.351 lineage, no significant alterations were observed. Whereas, most of the RBM residues in AY.1 varaint displayed enhanced contacts (Figure S9). All the potent NTD targeting antibodies recognize the ‘NTD supersite’, consisting of the N1, N3 and N5 loop (Lok, 2021). In the wild-type structure, NTD displayed a total of 2,843 local contacts with 52% of the NTD residues having a degree of >10. The higher degree residues comprised 8 out of the 17 characteristic mutation sites in NTD across the key lineages (T19, H69, Y144, E156, R158, R190, R246 & D253). Although B.1.1.7 intra-NTD network is supported by 233 residues with higher degree contacts, three deletions in the N2 loop of the NTD (H69del, V70del & Y144del) greatly perturb the local interaction network which was observed to mediate around 10, 9, 15 local contacts each in the WT spike, respectively. Similar alterations were observed around furin cleavage site with 44% more number of contacts within protein residues.

These findings suggest that these combinations of mutation present in lineages perturb the local interactions other than exact point of substitution and we further speculate that these rewired contacts may influence functionally important regions.

### Molecular dynamics simulations of full-length D614G spike showed structurally flexible regions

Collective internal motions in proteins are linked with the full-length spike structural model of wild-type and one mutant structure of spike protein in an open state. From here the 1-RBD-up spike structures bound to hACE2 were inserted into the membrane and each system was simulated for 450 ns each (Figure 6A). In comparison to the starting structure, root mean square deviations were plotted for each domain (Figure 6B). Our analysis revealed increased stability of the D614G protein (0.89 ± 0.007 nm) as compared to the WT spike system (Figure S10). Interestingly, a relative decrease in the deviation of mutant NTD (0.63 ± 0.002 nm) and RBD regions (0.74 ± 0.005 nm) was found, as shown in Figure 6B. In addition to RBD and NTD, high flexibility was found in the HR2 and CTD which have been linked to post fusion event (Huang et al., 2020; Turoňová et al., 2020). Followed by the RBD, the SD1 (542-591 a.a.) and SD2 (592-681 a.a.) regions showed a similar trend in both the WT and D614G systems with an average RMSD of 0.61 ± 0.003 nm. Figure 6C shows the representive snapshots highlighting dynamical changes within the NTD and RBD regions at different time frames. Specifically, the RBM in D614G (0.72 ± 0.002 nm) is observed to be relatively stable as compared to that of the WT system (1.00 ± 0.009 nm). Interestingly, each of the spike protein monomers is inter-twinned in such a fashion that it orients the RBD closer to the next chain NTD. We analysed the correlation between the RBD and NTD of each chain and our analyses revealed that the distances between these domains are higher in the wild-type (6.20 nm and 6.48 nm). In D614G trajectory, it corresponded to a relatively compact state with significantly less distance between the NTD and RBD (5.97 nm and 5.82 nm, respectively) as shown in Figure S11. This results in a compact conformation of the D614G and more importantly results in higher proximity with the hACE2 structure. Mapping of the flexibility parameter, root mean square fluctuations (RMSF) of both domains revealed significant stability in mutant structures (Figure S12). In wild-type, we observed distinct patches within the NTD (142-152, 180-184, and 248-257) and RBD (474 to 490, 496 to 503) that are highly mobile. Approximately 43.6% and 12.67% of the total mutations of the spike are present in the NTD and RBD domain, respectively. We speculate that these flexible regions evolve faster and hence play a key role in protein function.

**Figure 6:**
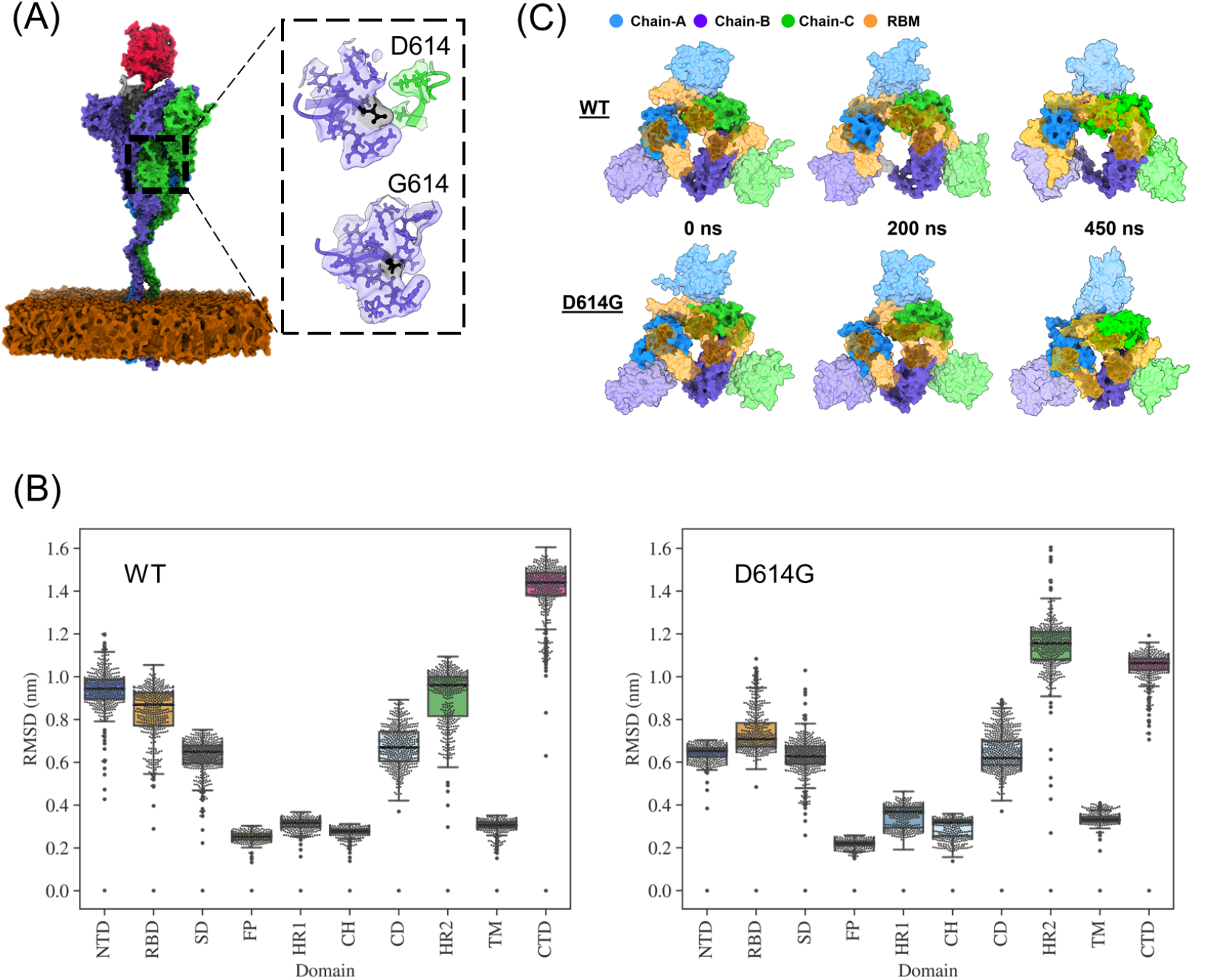
Dynamics of the wild-type and D614G mutant spike. (A) Full-length 1-RBD-up spike model in the POPC membrane bound to hACE2 is depicted. The zoomed-in image highlights the residues within 5 Å of the 614th residue in WT and mutant, where 614 is highlighted in black color. (B) Box plot showing the domain-wise distribution of RMSD in WT & mutant trajectories in the left and right panel, respectively. (C) Snapshots showing the dynamics of RBM with respect to the RBD and NTD in WT & D614G mutant spike protein as a function of time. The structures are shown in surface representation with chains A, B, and C colored in blue, purple, and green, respectively while brown shows the RBM motif.

## Discussion and conclusions

There are several driving forces for and against the selection determining the fitness advantage. The timely tracking of the sequencing of SARS-CoV-2 genomes in parallel with a large number of structural data has enabled several quick biological discoveries (Neher et al., 2020).During the ongoing pandemic, the genomes served as quick indicator maps and provided real-time tracking of key mutations (Rambaut et al., 2020; Faria et al., 2021). Especially, mutations on the SARS-CoV-2 spike protein are of substantial interest as they directly induces antibody response. Several vaccine candidates are also focused on spike protein (Martínez-Flores et al., 2021). Therefore, comprehensive efforts are required to decompose structural factors responsible for altered interactions in spike variants.

Here, we dynamically tracked genomic variations of SARS-CoV-2 spike protein submitted in GISAID over a span of 17 months which enabled a pan-spike mutational analysis that revealed more than 2500 mutations on the protein. We found precise mutational hotspots that are accruing more mutations and emphasize that there exists selective pressure on these stretches. Apart from well-characterised RBD region, several loops of NTD showed hypervariable regions. In addition, SARS-CoV-2 lineages mutations exhibit NTD mutations co-occurring with RBD, Linker and S2 domain, indicating coaxial framework of mutations. Further structure-based analysis of VOCs revealed that mutated side chains appear on surfaces and loops. Dense local networks around functionally important regions were significantly perturbed in VOCs.A caveat of the present structural analysis of VOCs is estimation of structural properties based on minimised structural snapshots. Extensive dynamic characterisation is provided for one mutated sidechain, D614G where we observed compactness in the NTD-RBD segments with D614G relatively stable. In our earlier detailed work on Delta variant surge in Delhi, we speculated that three locations: RBD, NTD, and furin cleavage site are collectively evolving for higher transmission (Dhar et al., 2021). Clearly, much remains to be learned, in terms of which mutational combinations and how their functional interactions (if any) change the biological function of the virus. Either way, analysis of high-throughput data and association with clinical markers must go hand in hand.

## Materials and Methods

### Genome data acquisition and analysis

In total, 1,624,465 spike protein sequences of SARS-CoV-2 genome were downloaded from the GISAID database submitted till 31 May, 2021 (Shu and McCauley, 2017). The low-quality sequences were removed and the genomes having high coverage greater than 90% were considered. 527,988 genomes passed the filtering criteria for specific seven geographies and were taken for further analysis. The additional details associated with these genomes including geography and date of submission was also considered for this study. Specifically, we selected 7 geographic regions, China (1,456), India (16,768), Brazil (3,936), South Africa (3,691), Australia (13,819), USA (287,843), and UK (200,475); constituting 62.24% of the total genomes. In addition, we also included 8,477 Indian genomes sequenced by NCDC, Delhi submitted to GISAID later to 31 May 2021. Any sequences without proper submission details were dropped out. The variant annotation was done using CoVsurver enabled by GISAID in GISAID’s EpiCoV database and Wuhan 2019-nCov genome (GISAID Accession ID: EPI ISL 402124) was used as the reference genome for mutational analysis (Okada et al., 2020). The quality check was performed and all the abrupt genomic sequences with more than 20 mutations per sample were removed. We also characterised month-wise genomic information to capture the time progression of spike variants. To remove the noise, we analysed mutations present in at least 1% of the total samples in selected time duration. We assigned lineage information for each sequence using Phylogenetic Assignment of Named Global Outbreak Lineages (PANGOLIN) resource (O’Toole et al., 2021). For analysing the inter-domain co-occurring combinations, we extracted the defining mutations for the six major lineages.

### Generation of hACE2-bound 1-RBD-up spike conformation

For extensive characterisation of spike protein using molecular dynamics simulations, we generated the full-length model of spike consisting of 1273 residues in an open conformation. The cryo-EM structure of the spike RBD-up state (PDB ID 6VSB) was taken as the template. In addition, the HR2 region was modeled using the NMR structure of the HR2 domain from SARS-CoV (PDB ID 2FXP) while that of the TM region was generated using the NMR structure of HIV-1 Env polyprotein (PDB ID 5JYN) having a homology of 100% and 29% with the spike protein, respectively. The PDB ID 5JYN was taken as it was the only template available for the TM region at the time of model generation. The missing residues in the structure were modelled as loops except for residues 1148-1161 which were restraint to form an alpha-helix. The structural model was generated in 3 segments: 14-1147, 1148-1213, and 1214-1273 and were joined together using the standard peptide-bond criteria in Chimera (Pettersen et al., 2004a). To obtain a complete atomistic model of hACE2, we took the X-ray structure of human ACE2 bound to the RBD domain (PDB ID: 6M0J) as template. The D614G mutation was then introduced in all three chains of the model in UCSF Chimera. The full-length spike model of 1-RBD-up state bound to hACE2 was oriented in the membrane containing 2068 POPC lipids. The membrane was generated using VMD and the spike structure was embedded in the membrane using InflateGRO2 (Humphrey et al., 1996; Kandt et al., 2007). The transmembrane residues (1214 to 1273) were positioned in the membrane by translating the protein along the Z-axis.

### Generation of wild-type and variant glycosylated spike protein models

The protein structure PDB ID 7A94 was taken as a reference structure to generate wild-type spike model. This structure is a higher resolution structure available for open bound conformation of spike consisting of 1146 residues (Benton et al., 2020). The complex structure depicts the 1-RBD-up conformation in spike protein, with the limited missing regions [71-75, 625-632, 677-688, 828-852, and 941-943] modeled using Modeller. The hACE2 structure in the template also had six residues missing patch that were added by taking PDB ID 1R42 as a reference. The complete 1146 spike-hACE2 structural model was used as a template to create variant models using Dunbrack 2010 backbone-dependent rotamer library in Chimera (Shapovalov and Dunbrack Jr, 2011). Further processing of steps was done using Gromacs toolkit. The deletions for B.1.1.7, B.1.617.2 and AY.1 spike were introduced and adjacent residues were joined by introducing the peptide bond. The mutated structural models were subjected to minimization in vacuum for extensive sidechain minimisation. The final step involved addition of glycosylation using the Glycan Reader & Modeler of Charmm-GUI (Jo et al., 2011).The detailed information of the site and type of glycosylation for each chain of spike was taken from Casalino et. al,2020 study(Casalino et al., 2020). All the glycosylated structural models are available on the github link of this paper.

### Analysis of structural properties

The relative solvent-accessible surface area for each residue in spike WT as well as the variants was calculated by Naccess (Hubbard and Thornton, 1993) and the secondary structure assessment was performed using STRIDE (Heinig and Frishman, 2004). We utilized network approach to study the alterations in the local contact network around the characteristic mutation sites. To obtain hydrophobic contacts, salt bridges and hydrogen bonding pattern for each of the structural model, the intra-protein interactions analysis has been carried out by using Pyinteraph and the interaction plotter (pymol plugin) was used to map the interactions identified by Pyinteraph on the 3D-spike structure (Tiberti et al., 2014). For the contact based intra-protein network, we took a cut-off of 3.5 Å to define the existence of a hydrogen bond between sidechain of residues and mainchain-sidechain residue atoms. For salt bridges, the charged residues were considered interacting if at least one pair of atoms is at a distance within the cut-off.

In addition, we also analysed inter-protein contacts between hACE2 and spike RBD region. We performed the network construction for our glycosylated spike structural models of WT and variants to capture the changes occurring within the binding interface. For this we considered the interface residues of RBD (400-508 a.a) and hACE2 (19-100 a.a, 300-360 a.a) and the co-ordinates for hACE2 were saved from the PDB ID 7A94. RINalyzer plugin in Cytoscape 3.8.2 was used to construct the networks (Demchak et al., 2014; Doncheva et al., 2011). The nodes represent the residues and the interactions between residues are edges of the network. A cut-off distance of 4Å was used to define a contact.

### Molecular Dynamics Simulations

The full-length hACE2-bound 1-RBD-up complex structure in membrane was placed in a rectangular box large enough to contain the protein and the membrane. Na+ ions were added to neutralize and TIP3P water representation was used to solvate the system (Mark and Nilsson, 2001). The simulations were carried out using GROMACS version 2018.3 by employing CHARMM36 all-atom force field (Abraham et al., 2015). Periodic boundary conditions were used and 1.2 nm was set as real space cut-off distance. Particle Mesh Ewald (PME) summation using the grid spacing of 0.16 nm was used combining with a fourth-order cubic interpolation to deduce the forces and potential in-between grid points (Darden et al., 1993). The Van der Waals cut-off was set to 1.2 nm. A 2 fs time step for numerical integration of the equations of motion was used and the coordinates were saved at every 100 ps. The initial systems were subjected to energy minimization using the steepest descent method. Temperature and pressure were maintained at 310 K and 1 bar using Nose-Hoover thermostat and Parrinello-Rahman barostat, respectively (Bussi et al., 2007; Parrinello and Rahman, 1981). Simulations of both wild-type and D614G systems were carried out for 450 ns each. All the molecular images were rendered using UCSF Chimera and ChimeraX (Pettersen et al., 2004b, 2021), VMD (Humphrey et al., 1996) and PyMOL (Schrödinger, LLC, 2015). The graphs and plots were generated using Python libraries.

## Acknowledgements

This work was supported by funding from CSIR and the Department of Science and Technology (DST). S.F. is supported by DST Early Career Award, S.R. by DST-Inspire fellowship, D.G. by CSIR-NET fellowship, and N.J. by DST-SERB fellowship. We acknowledge the support from CSIR-IGIB for infrastructure and CSIR-4PI for supercomputing facilities.

## Authors Contribution

All authors contributed to the study conception and design. SF and SR collected and analysed the structural and genomic data. MD simulation study and structural analysis were also performed by SR and SF. NB and DG contributed to the additional analyses and interpretation of the data. AP and MM analyzed the geographical distribution of mutations. LT supervised the study and helped in the interpretation of results. SF, SR and LT wrote the manuscript.

## Data availability

The data files related to the genomic analyses, structural models and MD simulation trajectories have been made available at https://github.com/CSB-Thukral-Lab/spike_genomic_and_structural_work.

## References

Abraham, M.J., Murtola, T., Schulz, R., Páll, S., Smith, J.C., Hess, B., Lindahl, E., 2015. GROMACS: High performance molecular simulations through multi-level parallelism from laptops to supercomputers. SoftwareX 1-2, 19–25. doi:10.1016/j.softx.2015.06.001.

Amanat, F., Thapa, M., Lei, T., Ahmed, S.M.S., Adelsberg, D.C., Carreño, J.M., Strohmeier, S., Schmitz, A.J., Zafar, S., Zhou, J.Q., et al., 2021. Sars-cov-2 mrna vaccination induces functionally diverse antibodies to ntd, rbd, and s2. Cell 184, 3936–3948.

Andersen, K.G., Rambaut, A., Lipkin, W.I., Holmes, E.C., Garry, R.F., 2020. The proximal origin of sars-cov-2. Nature medicine 26, 450–452.

Annavajhala, M.K., Mohri, H., Wang, P., Nair, M., Zucker, J.E., Sheng, Z., Gomez-Simmonds, A., Kelley, A.L., Tagliavia, M., Huang, Y., et al., 2021. A novel and expanding sars-cov-2 variant, b. 1.526, identified in new york. medRxiv.

Barnes, C.O., Jette, C.A., Abernathy, M.E., Dam, K.M.A., Esswein, S.R., Gristick, H.B., Malyutin, A.G., Sharaf, N.G., Huey-Tubman, K.E., Lee, Y.E., Robbiani, D.F., Nussenzweig, M.C., West, A.P., Bjorkman, P.J., 2020. SARS-CoV-2 neutralizing antibody structures inform therapeutic strategies. Nature 588, 682–687. doi:10.1038/s41586-020-2852-1.

Benton, D.J., Wrobel, A.G., Xu, P., Roustan, C., Martin, S.R., Rosenthal, P.B., Skehel, J.J., Gamblin, S.J., 2020. Receptor binding and priming of the spike protein of SARS-CoV-2 for membrane fusion. Nature 588, 327–330. doi:10.1038/s41586-020-2772-0.

Bussi, G., Donadio, D., Parrinello, M., 2007. Canonical sampling through velocity rescaling. The Journal of Chemical Physics 126, 014101. doi:10.1063/1.2408420.

Casalino, L., Gaieb, Z., Goldsmith, J.A., Hjorth, C.K., Dommer, A.C., Harbison, A.M., Fogarty, C.A., Barros, E.P., Taylor, B.C., McLellan, J.S., Fadda, E., Amaro, R.E., 2020. Beyond shielding: The roles of glycans in the SARS-CoV-2 spike protein. ACS Central Science 6, 1722–1734. doi:10.1021/acscentsci.0c01056.

Chen, X., Li, R., Pan, Z., Qian, C., Yang, Y., You, R., Zhao, J., Liu, P., Gao, L., Li, Z., Huang, Q., Xu, L., Tang, J., Tian, Q., Yao, W., Hu, L., Yan, X., Zhou, X., Wu, Y., Deng, K., Zhang, Z., Qian, Z., Chen, Y., Ye, L., 2020. Human monoclonal antibodies block the binding of SARS-CoV-2 spike protein to angiotensin converting enzyme 2 receptor. Cellular & Molecular Immunology 17, 647–649. doi:10.1038/s41423-020-0426-7.

Chi, X., Yan, R., Zhang, J., Zhang, G., Zhang, Y., Hao, M., Zhang, Z., Fan, P., Dong, Y., Yang, Y., Chen, Z., Guo, Y., Zhang, J., Li, Y., Song, X., Chen, Y., Xia, L., Fu, L., Hou, L., Xu, J., Yu, C., Li, J., Zhou, Q., Chen, W., 2020. A neutralizing human antibody binds to the n-terminal domain of the spike protein of SARS-CoV-2. Science 369, 650–655. doi:10.1126/science.abc6952.

Dai, L., Gao, G.F., 2021. Viral targets for vaccines against covid-19. Nature Reviews Immunology 21, 73–82.

Darden, T., York, D., Pedersen, L., 1993. Particle mesh ewald: AnN·log(n) method for ewald sums in large systems. The Journal of Chemical Physics 98, 10089–10092. doi:10.1063/1.464397.

Demchak, B., Hull, T., Reich, M., Liefeld, T., Smoot, M., Ideker, T., Mesirov, J.P., 2014. Cytoscape: the network visualization tool for genomespace workflows. F1000Research 3.

Dhar, M.S., Marwal, R., Radhakrishnan, V., Ponnusamy, K., Jolly, B., Bhoyar, R.C., Fatihi, S., Datta, M., Singh, P., Sharma, U., Ujjainia, R., Naushin, S., Bhateja, N., Divakar, M.K., Sardana, V., Singh, M.K., Imran, M., Senthivel, V., Maurya, R., Jha, N., Mehta, P., Rophina, M., Arvinden, V., Chaudhary, U., Thukral, L., Pandey, R., Dash, D., Faruq, M., Lall, H., Gogia, H., Madan, P., Kulkarni, S., Chauhan, H., Sengupta, S., Kabra, S., (INSACOG), T.I.S.C..G.C., Singh, S.K., Agrawal, A., Rakshit, P., 2021. Genomic characterization and epidemiology of an emerging sars-cov-2 variant in delhi, india. medRxiv doi:10.1101/2021.06.02.21258076.

Doncheva, N.T., Klein, K., Domingues, F.S., Albrecht, M., 2011. Analyzing and visualizing residue networks of protein structures. Trends in biochemical sciences 36, 179–182.

Faria, N.R., Mellan, T.A., Whittaker, C., Claro, I.M., Candido, D.d.S., Mishra, S., Crispim, M.A.E., Sales, F.C.S., Hawryluk, I., McCrone, J.T., Hulswit, R.J.G., Franco, L.A.M., Ramundo, M.S., de Jesus, J.G., Andrade, P.S., Coletti, T.M., Ferreira, G.M., Silva, C.A.M., Manuli, E.R., Pereira, R.H.M., Peixoto, P.S., Kraemer, M.U.G., Gaburo, N., Camilo, C.d.C., Hoeltgebaum, H., Souza, W.M., Rocha, E.C., de Souza, L.M., de Pinho, M.C., Araujo, L.J.T., Malta, F.S.V., de Lima, A.B., Silva, J.d.P., Zauli, D.A.G., Ferreira, A.C.d.S., Schnekenberg, R.P., Laydon, D.J., Walker, P.G.T., Schlüter, H.M., dos Santos, A.L.P., Vidal, M.S., Del Caro, V.S., Filho, R.M.F., dos Santos, H.M., Aguiar, R.S., Proença-Modena, J.L., Nelson, B., Hay, J.A., Monod, M., Miscouridou, X., Coupland, H., Sonabend, R., Vollmer, M., Gandy, A., Prete, C.A., Nascimento, V.H., Suchard, M.A., Bowden, T.A., Pond, S.L.K., Wu, C.H., Ratmann, O., Ferguson, N.M., Dye, C., Loman, N.J., Lemey, P., Rambaut, A., Fraiji, N.A., Carvalho, M.d.P.S.S., Pybus, O.G., Flaxman, S., Bhatt, S., Sabino, E.C., 2021. Genomics and epidemiology of the p.1 sars-cov-2 lineage in manaus, brazil. Science 372, 815–821. doi:10.1126/science.abh2644.

Graham, C., Seow, J., Huettner, I., Khan, H., Kouphou, N., Acors, S., Winstone, H., Pickering, S., Galao, R.P., Lista, M.J., et al., 2021. Impact of the b. 1.1. 7 variant on neutralizing monoclonal antibodies recognizing diverse epitopes on sars–cov–2 spike. bioRxiv.

Groves, D.C., Rowland-Jones, S.L., Angyal, A., 2021. The d614g mutations in the sars-cov-2 spike protein: Implications for viral infectivity, disease severity and vaccine design. Biochemical and biophysical research communications 538, 104–107.

Heinig, M., Frishman, D., 2004. Stride: a web server for secondary structure assignment from known atomic coordinates of proteins. Nucleic acids research 32, W500–W502.

Hoffmann, M., Kleine-Weber, H., Pöhlmann, S., 2020a. A multibasic cleavage site in the spike protein of sars-cov-2 is essential for infection of human lung cells. Molecular cell 78, 779–784.

Hoffmann, M., Kleine-Weber, H., Schroeder, S., Krüger, N., Herrler, T., Erichsen, S., Schiergens, T.S., Herrler, G., Wu, N.H., Nitsche, A., Müller, M.A., Drosten, C., Pöhlmann, S., 2020b. SARS-CoV-2 cell entry depends on ACE2 and TMPRSS2 and is blocked by a clinically proven protease inhibitor. Cell 181, 271–280.e8. doi:10.1016/j.cell.2020.02.052.

Huang, Y., Yang, C., Xu, X.f., Xu, W., Liu, S.w., 2020. Structural and functional properties of sars-cov-2 spike protein: potential antivirus drug development for covid-19. Acta Pharmacologica Sinica 41, 1141–1149.

Hubbard, S.J., Thornton, J.M., 1993. naccess. Computer Program, Department of Biochemistry and Molecular Biology, University College London 2.

Humphrey, W., Dalke, A., Schulten, K., 1996. VMD: Visual molecular dynamics. Journal of Molecular Graphics 14, 33–38. doi:10.1016/0263-7855(96)00018-5.

Isabel, S., Graña-Miraglia, L., Gutierrez, J.M., Bundalovic-Torma, C., Groves, H.E., Isabel, M.R., Eshaghi, A., Patel, S.N., Gubbay, J.B., Poutanen, T., et al., 2020. Evolutionary and structural analyses of sars-cov-2 d614g spike protein mutation now documented worldwide. Scientific reports 10, 1–9.

Jo, S., Song, K.C., Desaire, H., MacKerell Jr, A.D., Im, W., 2011. Glycan reader: automated sugar identification and simulation preparation for carbohydrates and glycoproteins. Journal of computational chemistry 32, 3135–3141.

Ju, B., Zhang, Q., Ge, J., Wang, R., Sun, J., Ge, X., Yu, J., Shan, S., Zhou, B., Song, S., Tang, X., Yu, J., Lan, J., Yuan, J., Wang, H., Zhao, J., Zhang, S., Wang, Y., Shi, X., Liu, L., Zhao, J., Wang, X., Zhang, Z., Zhang, L., 2020. Human neutralizing antibodies elicited by SARS-CoV-2 infection. Nature 584, 115–119. doi:10.1038/s41586-020-2380-z.

Kandt, C., Ash, W.L., Tieleman, D.P., 2007. Setting up and running molecular dynamics simulations of membrane proteins. Methods 41, 475–488. doi:10.1016/j.ymeth.2006.08.006.

Kannan, S.R., Spratt, A.N., Cohen, A.R., Naqvi, S.H., Chand, H.S., Quinn, T.P., Lorson, C.L., Byrareddy, S.N., Singh, K., 2021. Evolutionary analysis of the delta and delta plus variants of the sars-cov-2 viruses. Journal of Autoimmunity, 102715.

Korber, B., Fischer, W.M., Gnanakaran, S., Yoon, H., Theiler, J., Abfalterer, W., Hengartner, N., Giorgi, E.E., Bhattacharya, T., Foley, B., Hastie, K.M., Parker, M.D., Partridge, D.G., Evans, C.M., Freeman, T.M., de Silva, T.I., McDanal, C., Perez, L.G., Tang, H., Moon-Walker, A., Whelan, S.P., LaBranche, C.C., Saphire, E.O., Montefiori, D.C., Angyal, A., Brown, R.L., Carrilero, L., Green, L.R., Groves, D.C., Johnson, K.J., Keeley, A.J., Lindsey, B.B., Parsons, P.J., Raza, M., Rowland-Jones, S., Smith, N., Tucker, R.M., Wang, D., Wyles, M.D., 2020. Tracking changes in SARS-CoV-2 spike: Evidence that d614g increases infectivity of the COVID-19 virus. Cell 182, 812–827.e19. doi:10.1016/j.cell.2020.06.043.

Lan, J., Ge, J., Yu, J., Shan, S., Zhou, H., Fan, S., Zhang, Q., Shi, X., Wang, Q., Zhang, L., Wang, X., 2020. Structure of the SARS-CoV-2 spike receptor-binding domain bound to the ACE2 receptor. Nature 581, 215–220. doi:10.1038/s41586-020-2180-5.

Liu, L., Wang, P., Nair, M.S., Yu, J., Rapp, M., Wang, Q., Luo, Y., Chan, J.F.W., Sahi, V., Figueroa, A., et al., 2020. Potent neutralizing antibodies against multiple epitopes on sars-cov-2 spike. Nature 584, 450–456.

Lok, S.M., 2021. An ntd supersite of attack. Cell host & microbe 29, 744–746.

Mark, P., Nilsson, L., 2001. Structure and dynamics of the TIP3p, SPC, and SPC/e water models at 298 k. The Journal of Physical Chemistry A 105, 9954–9960. doi:10.1021/jp003020w.

Martínez-Flores, D., Zepeda-Cervantes, J., Cruz-Reséndiz, A., Aguirre-Sampieri, S., Sampieri, A., Vaca, L., 2021. Sars-cov-2 vaccines based on the spike glycoprotein and implications of new viral variants. Frontiers in Immunology 12.

Mittal, A., Manjunath, K., Ranjan, R.K., Kaushik, S., Kumar, S., Verma, V., 2020. Covid-19 pandemic: Insights into structure, function, and hace2 receptor recognition by sars-cov-2. PLoS pathogens 16, e1008762.

Neher, R.A., Dyrdak, R., Druelle, V., Hodcroft, E.B., Albert, J., 2020. Potential impact of seasonal forcing on a SARS-CoV-2 pandemic. Swiss Med Wkly doi:10.1101/2020.02.13.20022806.

Okada, P., Buathong, R., Phuygun, S., Thanadachakul, T., Parnmen, S., Wongboot, W., Waicharoen, S., Wacharapluesadee, S., Uttayamakul, S., Vachiraphan, A., Chittaganpitch, M., Mekha, N., Janejai, N., Iamsirithaworn, S., Lee, R.T., Maurer-Stroh, S., 2020. Early transmission patterns of coronavirus disease 2019 (COVID-19) in travellers from wuhan to thailand, january 2020. Eurosurveillance 25. doi:10.2807/1560-7917.es.2020.25.8.2000097.

O’Toole, Á., Scher, E., Underwood, A., Jackson, B., Hill, V., McCrone, J.T., Colquhoun, R., Ruis, C., Abu-Dahab, K., Taylor, B., et al., 2021. Assignment of epidemiological lineages in an emerging pandemic using the pangolin tool. Virus Evolution 30.

Parrinello, M., Rahman, A., 1981. Polymorphic transitions in single crystals: A new molecular dynamics method. Journal of Applied Physics 52, 7182–7190. doi:10.1063/1.328693.

Pettersen, E.F., Goddard, T.D., Huang, C.C., Couch, G.S., Greenblatt, D.M., Meng, E.C., Ferrin, T.E., 2004a. UCSF chimera?a visualization system for exploratory research and analysis. Journal of Computational Chemistry 25, 1605–1612. doi:10.1002/jcc.20084.

Pettersen, E.F., Goddard, T.D., Huang, C.C., Couch, G.S., Greenblatt, D.M., Meng, E.C., Ferrin, T.E., 2004b. Ucsf chimera—a visualization system for exploratory research and analysis. Journal of computational chemistry 25, 1605–1612.

Pettersen, E.F., Goddard, T.D., Huang, C.C., Meng, E.C., Couch, G.S., Croll, T.I., Morris, J.H., Ferrin, T.E., 2021. Ucsf chimerax: Structure visualization for researchers, educators, and developers. Protein Science 30, 70–82.

Rambaut, A., Loman, N., Pybus, O., Barclay, W., Barrett, J., Carabelli, A., Connor, T., Peacock, T., Robertson, D.L., Volz, E., 2020. Preliminary genomic characterisation of an emergent sars-cov-2 lineage in the uk defined by a novel set of spike mutations.

Rees-Spear, C., Muir, L., Griffith, S., Heaney, J., Aldon, Y., Snitselaar, J., Thomas, P., Graham, C., Seow, J., Lee, N., Rosa, A., Roustan, C., Houlihan, C., Sanders, R., Gupta, R., Cherepanov, P., Stauss, H., Nastouli, E., Doores, K., van Gils, M., McCoy, L., 2021. The impact of spike mutations on SARS-CoV-2 neutralization. Cell Reports doi:10.1101/2021.01.15.426849.

Resende, P.C., Bezerra, J.F., Vasconcelos, R., Arantes, I., Appolinario, L., Mendonça, A.C., Paixao, A.C., Rodrigues, A.C.D., Silva, T., Rocha, A.S., et al., 2021. Spike e484k mutation in the first sars-cov-2 reinfection case confirmed in brazil, 2020. Virological [Internet] 10.

Schrödinger, LLC, 2015. The PyMOL molecular graphics system, version 1.8.

Shang, J., Ye, G., Shi, K., Wan, Y., Luo, C., Aihara, H., Geng, Q., Auerbach, A., Li, F., 2020. Structural basis of receptor recognition by sars-cov-2. Nature 581, 221–224.

Shapovalov, M.V., Dunbrack Jr, R.L., 2011. A smoothed backbone-dependent rotamer library for proteins derived from adaptive kernel density estimates and regressions. Structure 19, 844–858.

Shu, Y., McCauley, J., 2017. GISAID: Global initiative on sharing all influenza data – from vision to reality. Eurosurveillance 22. doi:10.2807/1560-7917.es.2017.22.13.30494.

Tegally, H., Wilkinson, E., Giovanetti, M., Iranzadeh, A., Fonseca, V., Giandhari, J., Doolabh, D., Pillay, S., San, E.J., Msomi, N., et al., 2020. Emergence and rapid spread of a new severe acute respiratory syndrome-related coronavirus 2 (sars-cov-2) lineage with multiple spike mutations in south africa. medRxiv.

Tegally, H., Wilkinson, E., Giovanetti, M., Iranzadeh, A., Fonseca, V., Giandhari, J., Doolabh, D., Pillay, S., San, E.J., Msomi, N., et al., 2021. Detection of a sars-cov-2 variant of concern in south africa. Nature 592, 438–443.

Tiberti, M., Invernizzi, G., Lambrughi, M., Inbar, Y., Schreiber, G., Papaleo, E., 2014. Pyinteraph: a framework for the analysis of interaction networks in structural ensembles of proteins. Journal of chemical information and modeling 54, 1537–1551.

Turoňová, B., Sikora, M., Schürmann, C., Hagen, W.J., Welsch, S., Blanc, F.E., von Bülow, S., Gecht, M., Bagola, K., Hörner, C., et al., 2020. In situ structural analysis of sars-cov-2 spike reveals flexibility mediated by three hinges. Science 370, 203–208.

Volz, E., Hill, V., McCrone, J.T., Price, A., Jorgensen, D., O’Toole, Á., Southgate, J., Johnson, R., Jackson, B., Nascimento, F.F., et al., 2021. Evaluating the effects of sars-cov-2 spike mutation d614g on transmissibility and pathogenicity. Cell 184, 64–75.

Walls, A.C., Park, Y.J., Tortorici, M.A., Wall, A., McGuire, A.T., Veesler, D., 2020. Structure, function, and antigenicity of the SARS-CoV-2 spike glycoprotein. Cell 181, 281–292.e6. doi:10.1016/j.cell.2020.02.058.

Wang, Y., Wang, L., Cao, H., Liu, C., 2021. Sars-cov-2 s1 is superior to the rbd as a covid-19 subunit vaccine antigen. Journal of medical virology 93, 892–898.

West Jr, A.P., Wertheim, J.O., Wang, J.C., Vasylyeva, T.I., Havens, J.L., Chowdhury, M.A., Gonzalez, E., Fang, C.E., Di Lonardo, S.S., Hughes, S., et al., 2021. Detection and characterization of the sars-cov-2 lineage b. 1.526 in new york. bioRxiv.

Xia, S., Lan, Q., Su, S., Wang, X., Xu, W., Liu, Z., Zhu, Y., Wang, Q., Lu, L., Jiang, S., 2020. The role of furin cleavage site in sars-cov-2 spike protein-mediated membrane fusion in the presence or absence of trypsin. Signal transduction and targeted therapy 5, 1–3.

Yurkovetskiy, L., Wang, X., Pascal, K.E., Tomkins-Tinch, C., Nyalile, T.P., Wang, Y., Baum, A., Diehl, W.E., Dauphin, A., Carbone, C., Veinotte, K., Egri, S.B., Schaffner, S.F., Lemieux, J.E., Munro, J.B., Rafique, A., Barve, A., Sabeti, P.C., Kyratsous, C.A., Dudkina, N.V., Shen, K., Luban, J., 2020. Structural and functional analysis of the d614g SARS-CoV-2 spike protein variant. Cell 183, 739–751.e8. doi:10.1016/j.cell.2020.09.032.

Zahradnik, J., Marciano, S., Shemesh, M., Zoler, E., Chiaravalli, J., Meyer, B., Dym, O., Elad, N., Schreiber, G., 2021. Sars-cov-2 rbd in vitro evolution follows contagious mutation spread, yet generates an able infection inhibitor. biorxiv.

Zhu, X., Mannar, D., Srivastava, S.S., Berezuk, A.M., Demers, J.P., Saville, J.W., Leopold, K., Li, W., Dimitrov, D.S., Tuttle, K.S., et al., 2021. Cryoelectron microscopy structures of the n501y sars-cov-2 spike protein in complex with ace2 and 2 potent neutralizing antibodies. PLoS biology 19, e3001237.

