## Supplementary material for "A rigorous framework for detecting SARS-CoV-2 spike protein mutational ensemble from genomic and structural features"

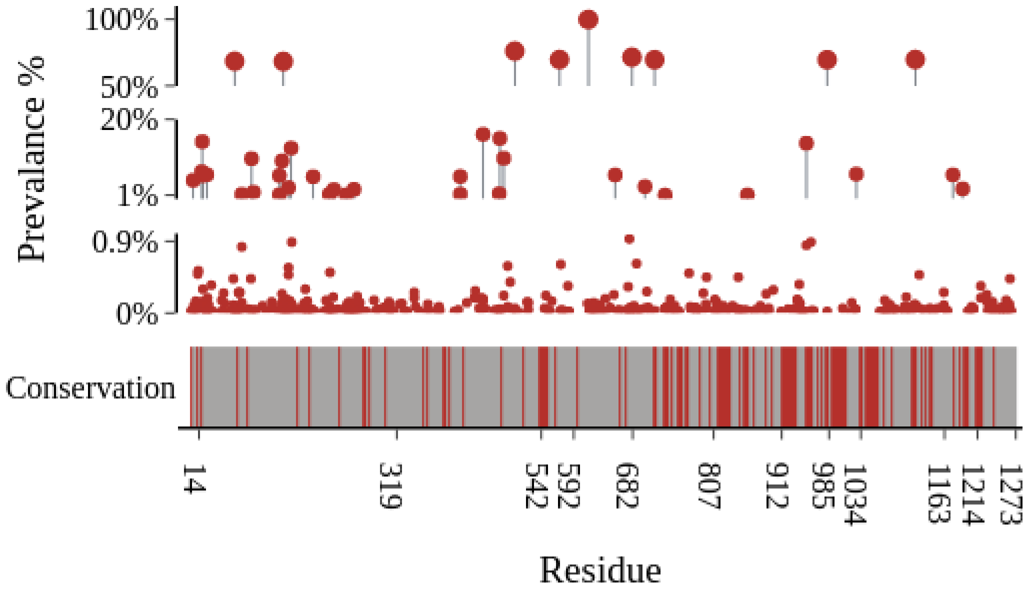


**Figure S1: Genomic variations in spike protein.** The needle plot showing the residue-wise global frequency of mutations in the spike protein, with prevalence of each mutation plotted against the Y-axis. On the basis of the prevalence, mutations are divided into three groups. The lower panel depicts the sequence conservation of spike protein residues with the boundaries of spike domains marked and residues having a conservation score =/> 7 are highlighted in red, whereas variable residues are in grey.

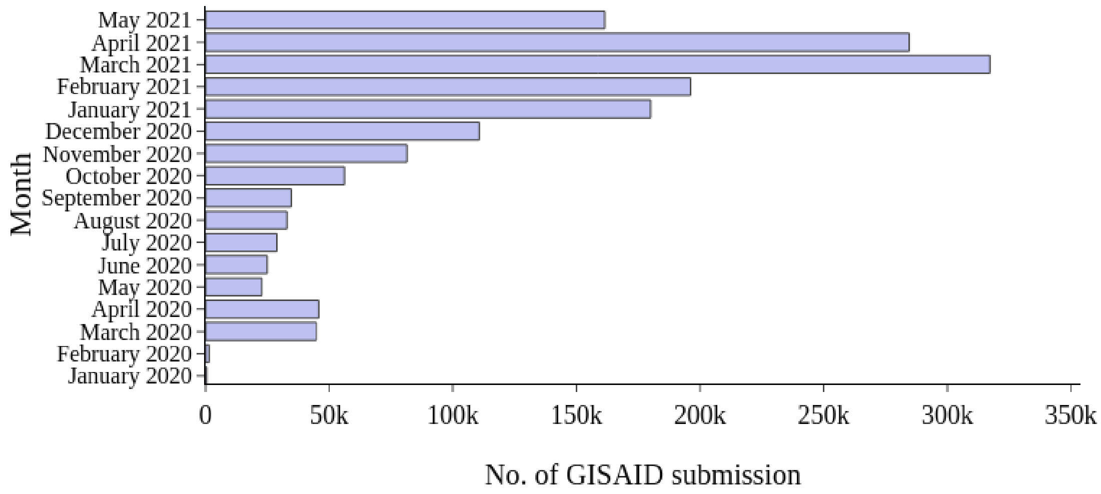


**Figure S2: Month-wise distribution of SARS-CoV-2 genomes.** The horizontal bar plot shows the distribution of the total no. of spike genomes submitted to GISAID in each month, since January 2020.

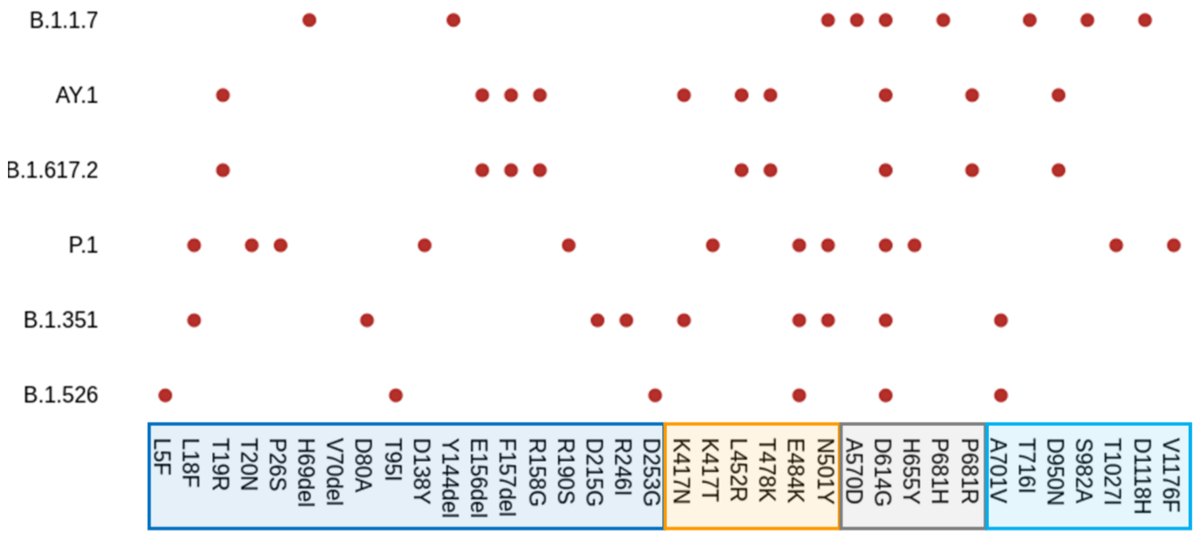


**Figure S3: Defining spike mutations for key lineages.** The scatter plot highlights the defining mutations present in each of these key lineages, where the NTD, RBD, Linker and S2 residues are marked in blue, yellow, grey and cyan, respectively.

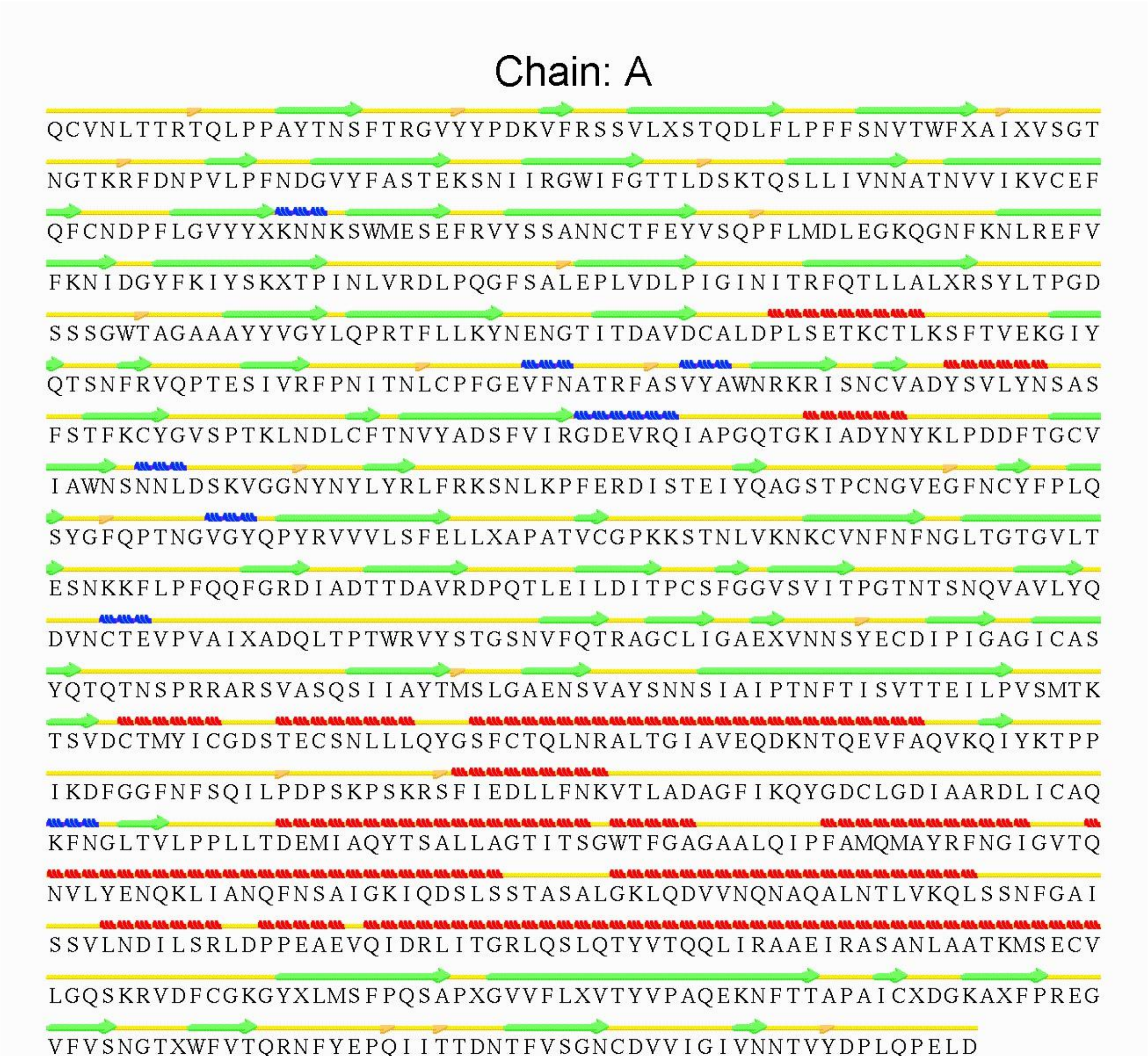


**Figure S4: Graphical summary for the secondary structural assessment of spike protein.** The secondary structural details for chain A (14-1146 a.a.) of the spike protein is shown where loops, helices, and sheets are displayed in yellow, green, and red respectively. $3_{10}$ helices are shown in blue.

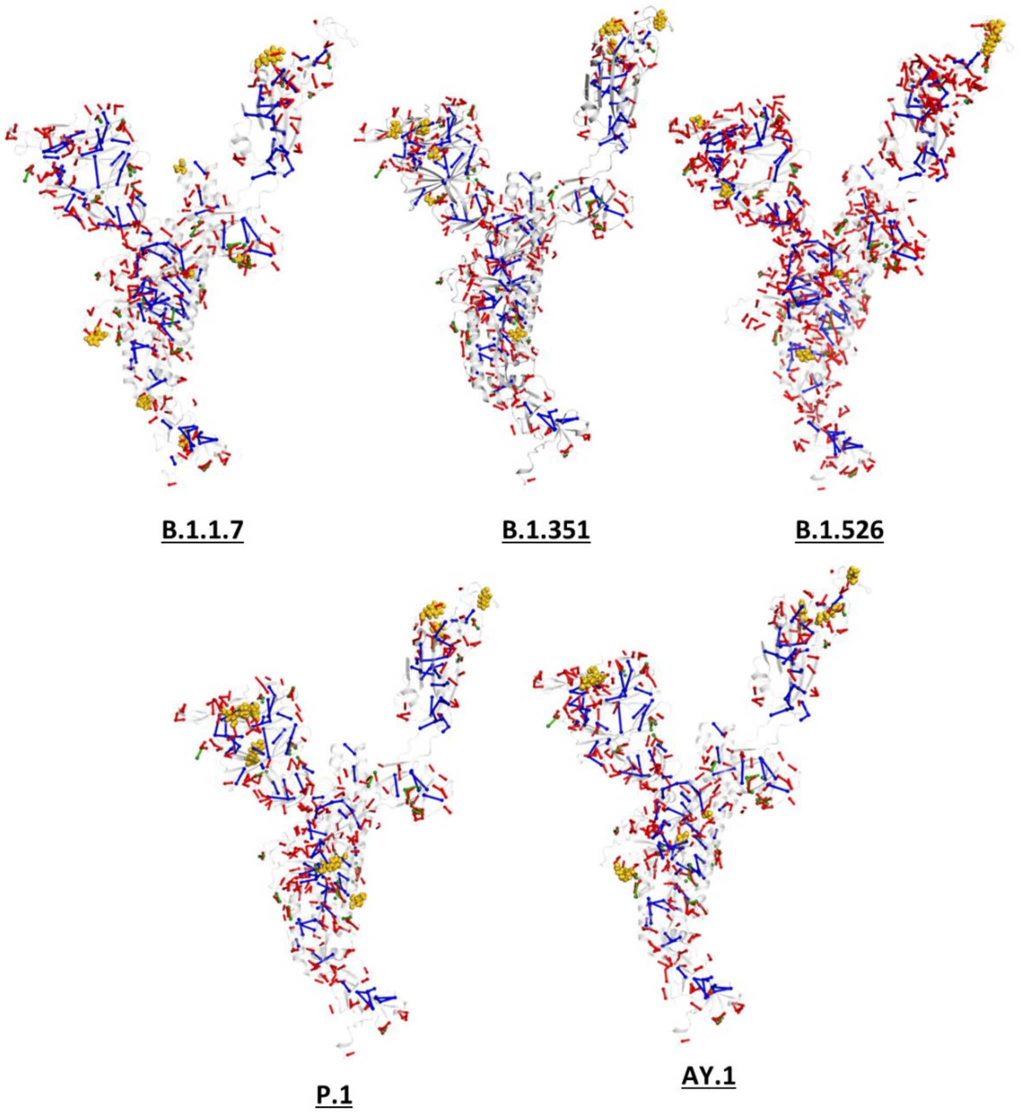


**Figure S5: Comparative global intramolecular network for the key spike variants.** The intramolecular interaction network is mapped on one of the chains for each of the spike variant, where the hydrogen bonds (unique to the variants), salt-bridges and hydrophobic interactions are shown in red, green, and blue, respectively. Mutations in each variant is highlighted in yellow.

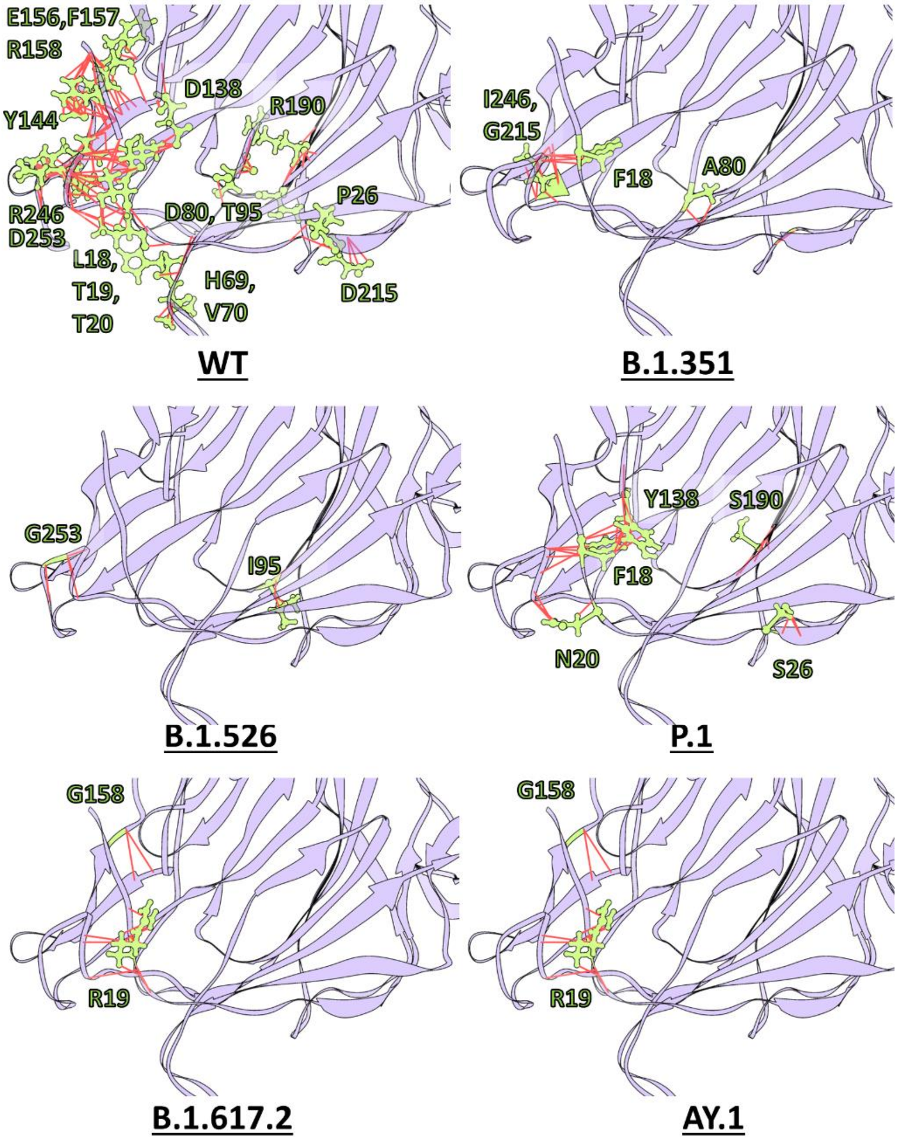


**Figure S6: Local intramolecular interactions mediated by the key mutations in the NTD.** The zoomed-in structural maps of the intramolecular interaction network around the mutation sites in the NTD of the key lineages are shown, where the mutations are highlighted in green ball-&-stick representation and their interactions within 4 Å are highlighted in red.

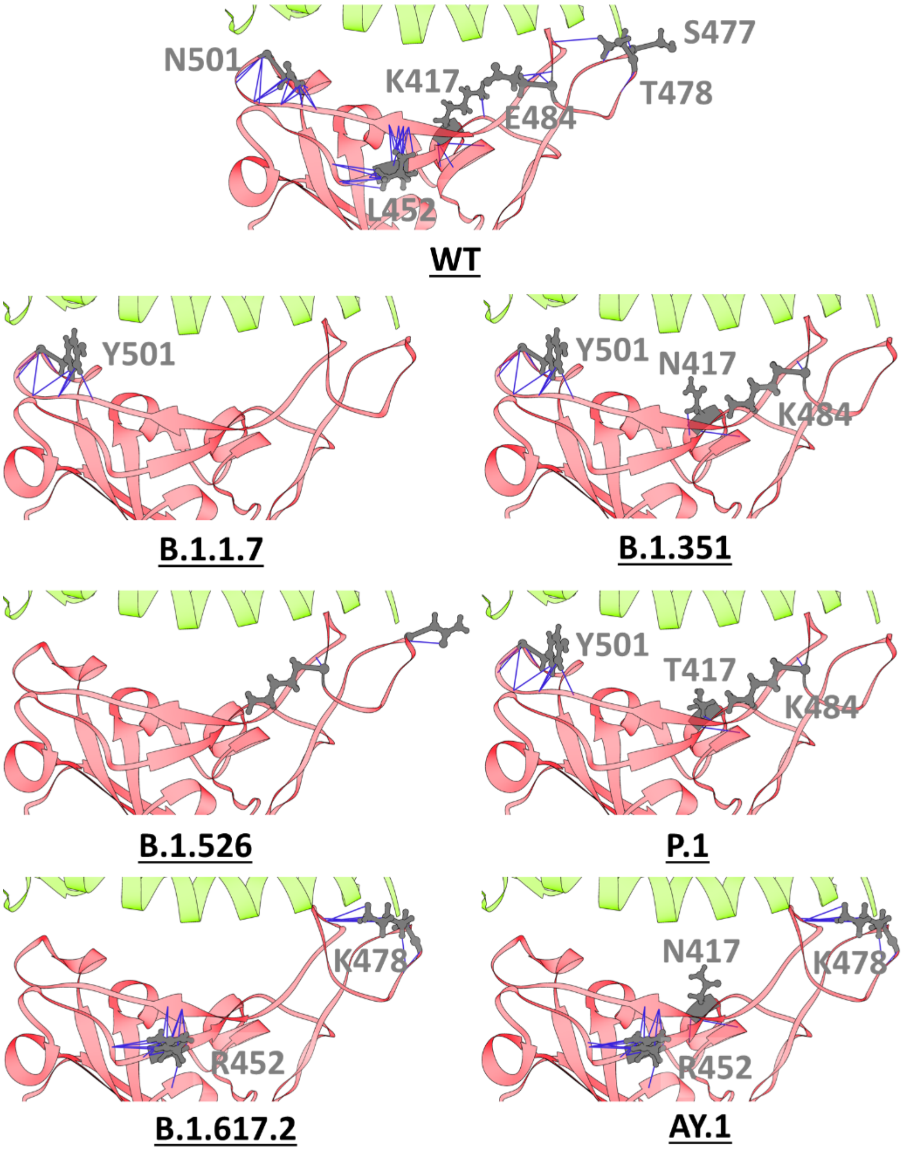


**Figure S7: Local intramolecular interactions mediated by the key mutations in the RBM**. The zoomed-in structural maps of the intramolecular interaction network around the mutation sites in the RBD of the key lineages are shown, where the mutations are highlighted in ball-&-stick representation and their interactions within 4 Å are highlighted in blue. Red and green ribbons represent RBM and hACE2, respectively.

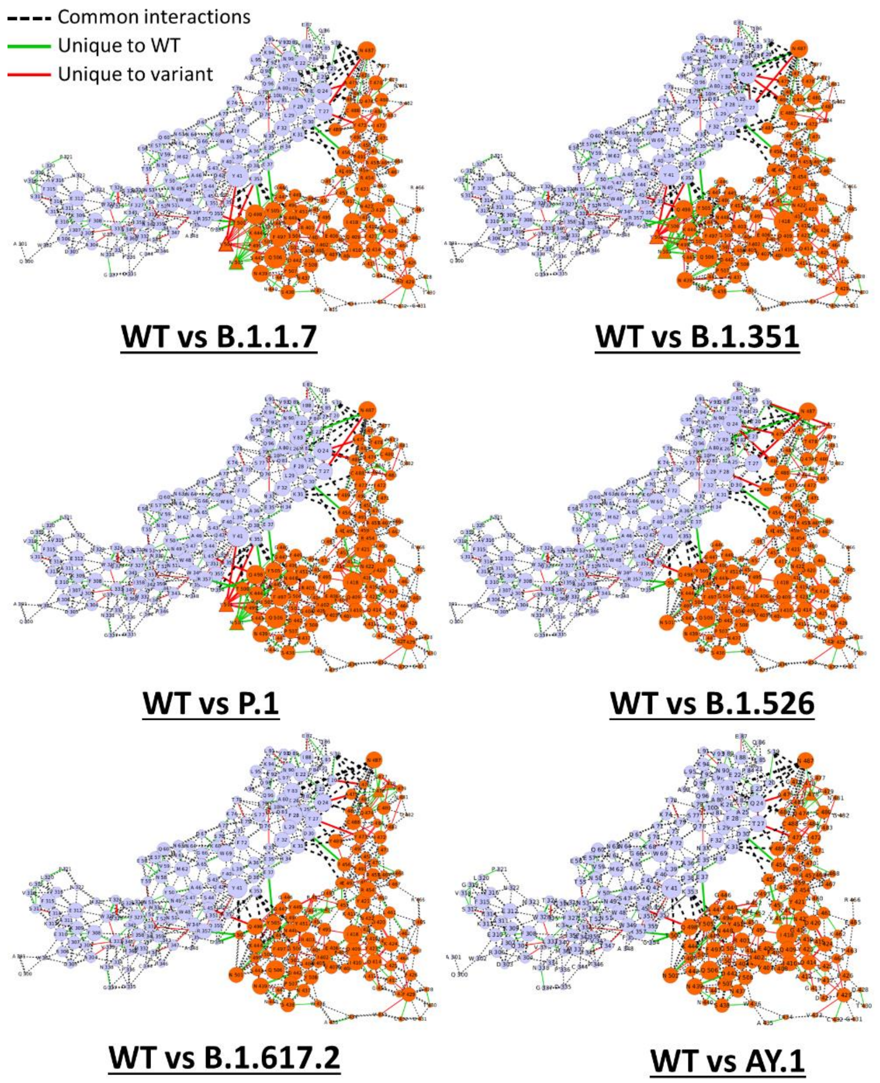


**Figure S8: Comparative global inter-molecular network for hACE2 bound RBD region in WT vs Variants.** Comparative inter-molecular interaction network for RBD-ACE2 in WT vs variant spike protein, where orange and light blue nodes represent the RBD and hACE2 residues, respectively and the degree of the nodes highlights the number of interactions mediated by each residue.

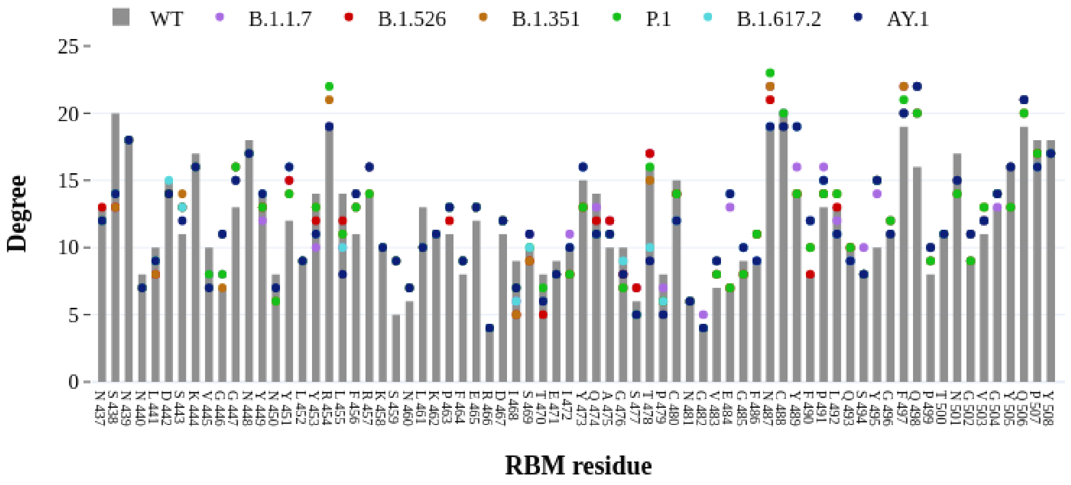


**Figure S9: Comparative distribution of degree for RBM residues.** The plot highlights the number of local interactions mediated by each residue of the RBM in the wild type and variants.

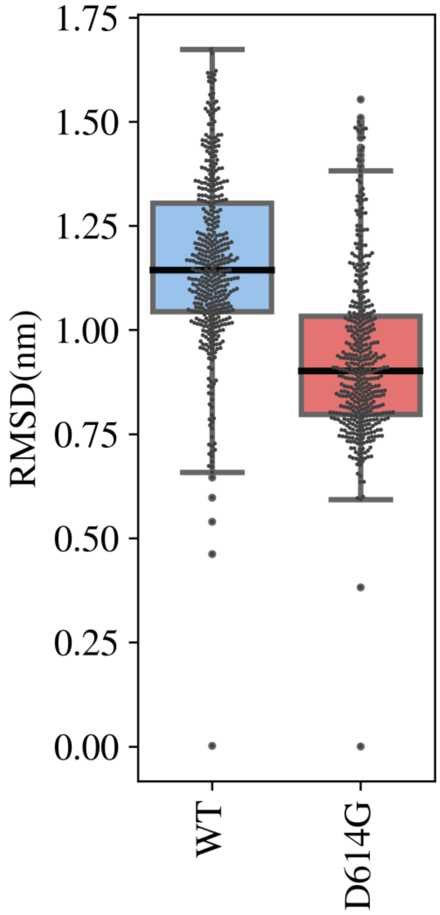


**Figure S10: Comparative dynamics of the wild and D614G-mutant trajectories.** Box plot showing the distribution of RMSD of the spike in wild and mutant trajectories of 1-RBD-up hACE2 bound spike system.

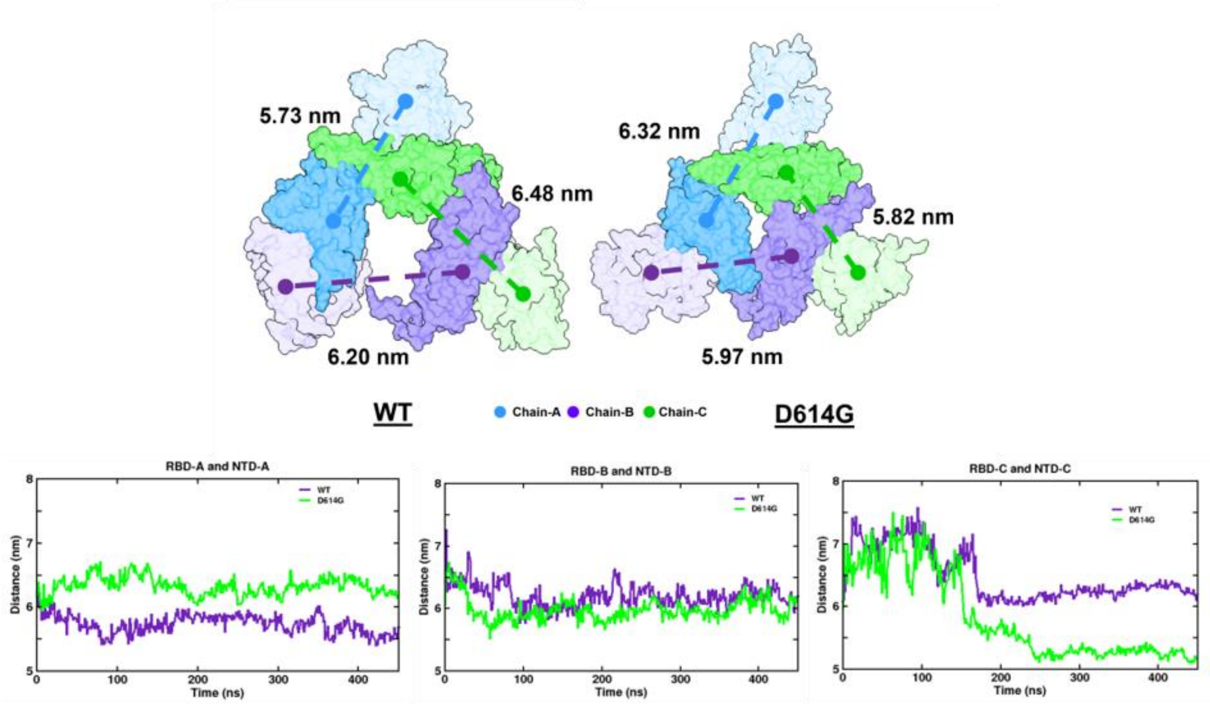


**Figure S11: NTD-RBD cross-talk in spike-wild and D614G.** The top-views of the wild-type and D614G NTD (high transparency) and RBD (low transparency) for each chain are shown in the snapshots, where chain A, B, and C are colored in blue, purple, and green, respectively. The average distance between the NTD and RBD of respective chains is represented with the dotted lines. The line plot shows the relative change in distances of the centre of mass of the NTD and RBD (within the chain) over 450 ns in wild-type and D614G mutant.

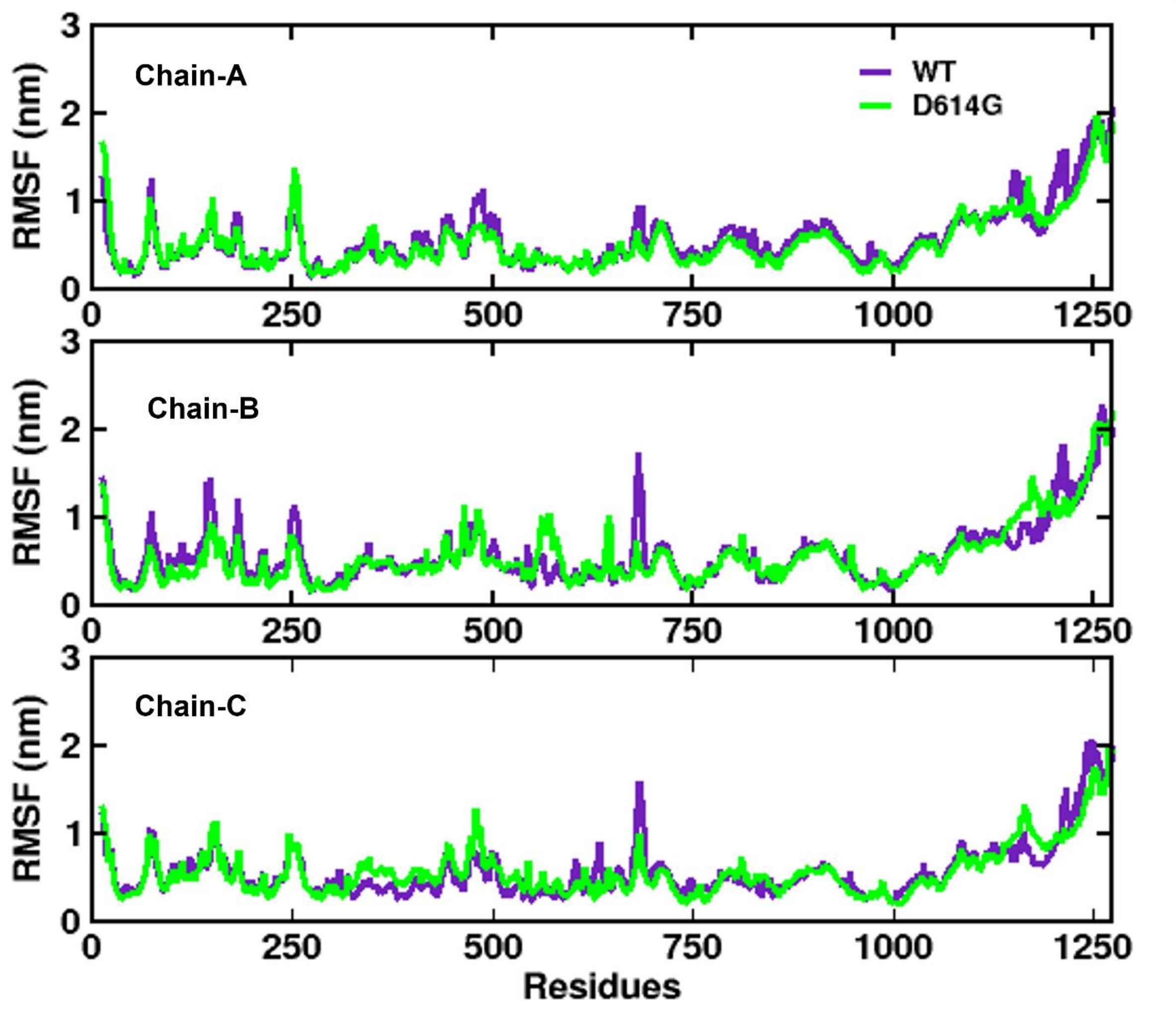


**Figure S12: Residue wise fluctuations across wild and D614G mutant trajectories.** Chain-wise root mean square fluctuation (RMSF) of spike protein in wild and mutant trajectories, where wild and D614G are colored in purple and green, respectively.
